# A *Candida glabrata* adhesin-like effector drives fitness and immunogenicity in the gut

**DOI:** 10.64898/2026.05.04.722752

**Authors:** Owen Jensen, Luke Hanson, Mathieu Hénault, Breanne E. Haskins, Emma Trujillo, Charlotte Brown, Tonya Brunetti, Maxwell McCabe, Brian C. Russo, Lydia Heasley, Kyla S. Ost

**Affiliations:** Department of Immunology and Microbiology, University of Colorado Anschutz Medical Campus, Aurora, Colorado, USA; Department of Biochemistry and Molecular Genetics, University of Colorado Anschutz Medical Campus, Aurora, Colorado, USA

**Keywords:** *Candida glabrata*, *Nakaseomyces glabratus*, immunoglobulin A, adhesins, colonization resistance, gut colonization

## Abstract

*Candida glabrata* is a leading cause of invasive candidiasis. The gut serves as its primary reservoir, yet factors governing colonization and pathogenic potential remain poorly defined. Here, we identify immunoglobulin A (IgA) as a key regulator of *C. glabrata* within the intestinal microbiome. We found that *C. glabrata* induces an IgA response in a strain-specific manner. Comparative transcriptional and proteomic analyses of IgA-inducing and non-inducing strains identified a putative adhesin, Awp11, whose expression correlated with IgA induction. Awp11 is directly targeted by IgA and is required for inducing *C. glabrata*-specific IgA and Th17 responses in vivo. Functionally, Awp11 promotes colonization of a complex intestinal microbiome, and intestinal IgA limits this advantage. In most strains, *AWP11* transcription is dynamic and limited by IgA in the gut. This identifies Awp11 as a key determinant of strain-dependent immunogenicity and gut colonization that *C. glabrata* may dynamically regulate to balance colonization and immune evasion.

## Introduction

*Candida* species asymptomatically colonize the gastrointestinal tract of 40-80% of healthy adults. While typically benign, these fungi are leading causes of opportunistic mucosal and invasive infections that collectively account for nearly 1 million deaths annually (1, 2). Recent studies have identified key interactions between the mucosal immune system and the intestinal microbiota that limit *Candida* colonization and damaging potential. However, these studies have overwhelmingly focused on a single species, *Candida albicans.* Non-*albicans* species are responsible for an increasing proportion of invasive candidiasis worldwide. Notably, *Candida glabrata* causes 20-40% of invasive Candidiasis, and is approaching or has recently surpassed *C. albicans* as the leading cause of invasive candidiasis in the United States and Western Europe (3, 4). Like *C. albicans, C. glabrata* asymptomatically colonizes the healthy human gut microbiota and can seed invasive infections from this reservoir (5, 6). Yet *C. glabrata* is genetically divergent from major *Candida* pathogens and has been recently reclassified as *Nakaseomyces glabratus* (though we will refer to it here as *C. glabrata*). Despite its clinical importance, the fungal effectors that support *C. glabrata* intestinal colonization, and the immune mechanisms regulating this fungus in the gut, remain poorly defined.

Immunoglobulin A (IgA) antibodies play a critical role in limiting the inflammatory and invasive potential of intestinal *Candida*. Studies on *C. albicans* have shown that IgA preferentially targets antigens expressed by the hyphal morphotype, and that colonization in the presence of IgA restricts hyphal growth and the expression of hyphae-associated virulence factors in the gut (7–9). In these same studies, *C. glabrata*-reactive IgA was detected in human intestinal wash and *C. glabrata* was shown to stimulate a species*-*specific IgA response in mice (9), suggesting a potential role for IgA in regulating *C. glabrata* in the gut.

Importantly, not all commensal fungi elicit IgA responses in the gut. Whereas *C. albicans* induces a robust, T cell-dependent IgA response and is heavily targeted by IgA during colonization, *Saccharomyces cerevisiae,* a nonpathogenic yeast commonly found in the human microbiome, does not (9). In *C. albicans*, IgA and CD4^+^ T cell responses depend on filamentation and the expression of hyphae-associated effectors, including the cytolytic toxin candidalysin and multiple adhesins (7, 9, 10). It was therefore unexpected that *C. glabrata*, a close genetic relative of *S. cerevisiae* that lacks the ability to form true hyphae, also stimulates a robust, antigen-specific IgA response in mice. This parallels observations in commensal bacteria, where IgA induction varies widely across species and strains (11). Although microbial phenotypes, such as the ability to adhere closely to the intestinal epithelium and access to gut-associated lymphoid tissue, have been linked to IgA induction (12, 13), the microbial determinants of IgA immunogenicity remain incompletely understood.

Here, we leverage a collection of clinical *C. glabrata* strains to define fungal mechanisms underlying strain-dependent IgA responses. We identify strain-specific expression of a signal putative adhesin, Awp11, as a key determinant of IgA induction. In one highly immunogenic strain, constitutively high *AWP11* expression is associated with a chromosomal translocation. Using this strain, we show that Awp11 is a dominant IgA-targeted antigen and is required for IgA induction and Th17 expansion. Functionally, we find that Awp11 promotes *C. glabrata* colonization of an intact microbiota, and that IgA ameliorates this advantage. Outside of this highly immunogenic strain, most strains dynamically regulate *AWP11* expression in response to the intestinal conditions. In one of these strains, *AWP11* expression is repressed in the context of an intestinal IgA response in the gut. Together, these findings identify Awp11 as central to immune and microbiota interactions with *C. glabrata,* both of which shaping the persistence of this opportunistic fungal pathogen in the gut.

## Results

### Anti-*C. glabrata* IgA responses are strain dependent

We previously demonstrated that the clinical *C. glabrata* strain Cg1, originally isolated from a human bloodstream infection, induces an intestinal IgA response in mice (9). However, it is unclear whether this response is universal across strains and how IgA regulates *C. glabrata* in the intestinal tract. To address this, we performed an in vivo screen of 34 clinical bloodstream isolates. Strains were grouped into five pools (5–7 strains per pool) and used to repeatedly colonize antibiotic-treated mice (Fig. 1A). Repeated gavage ensured continuous exposure despite strain-specific differences in colonization fitness; Cg5 was excluded because of poor growth in culture. A Cg1-only group served as a positive control. Fecal IgA reactivity to pooled strains grown in culture was assessed weekly by flow cytometry. No *C. glabrata*-reactive IgA was detected at day 7, but responses were detected by days 14–21 (Fig. S1A). Pool 4 induced the earliest and strongest IgA response, whereas pool 1 failed to induce detectable IgA at any time point. These kinetics indicate that *C. glabrata*-reactive IgA develops post-colonization rather than pre-existing in uncolonized mice.

**Figure 1.**
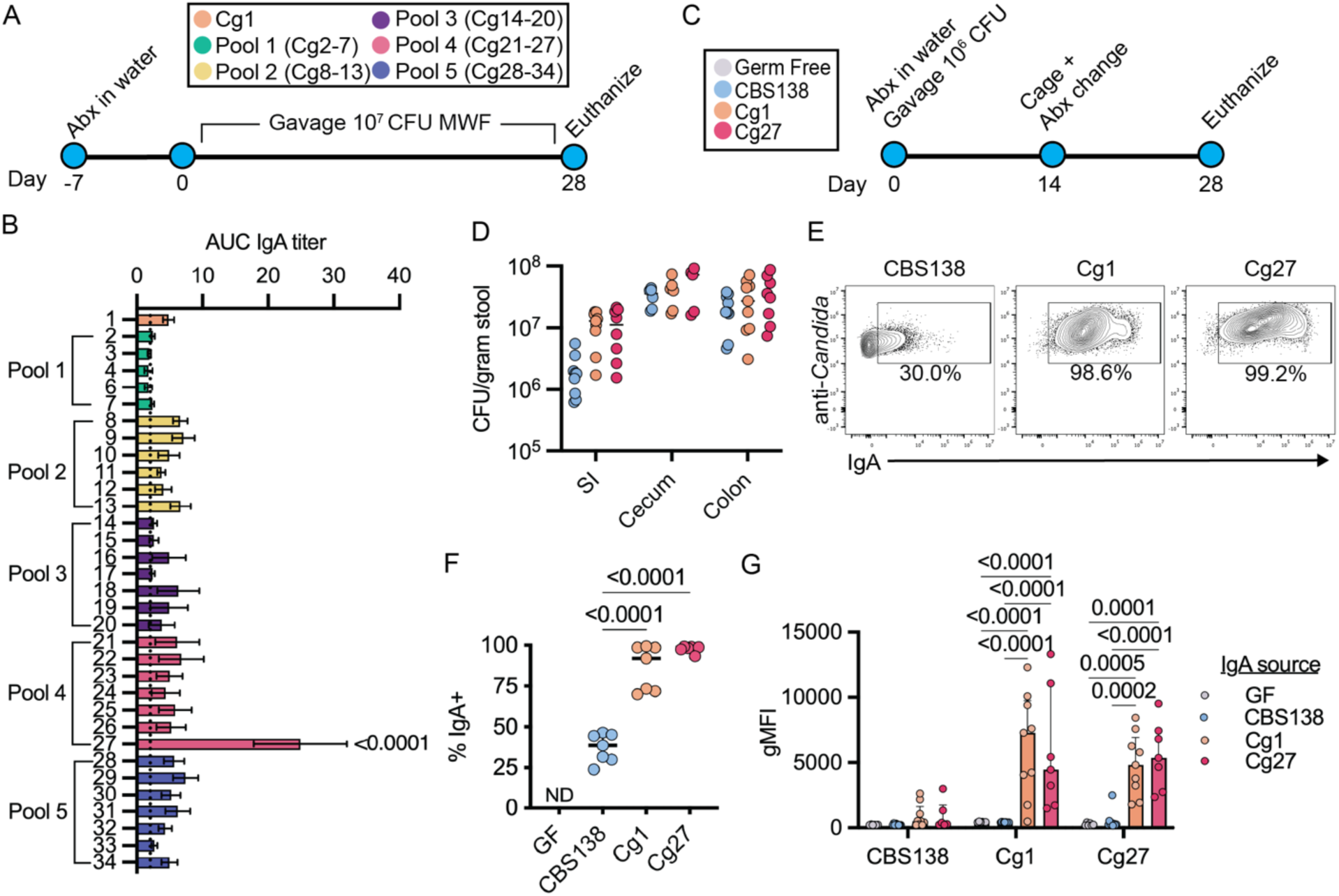
Anti-*C. glabrata* IgA responses are strain dependent. (A) Schematic of pooled screen for IgA-targeted *C. glabrata* isolates. (B) IgA binding from cleared intestinal wash to cultured strains from each strain pool. IgA reactivity quantified by flow cytometry and shown as area under the curve (AUC). (C) Schematic of GF mono-colonization experiment. (D) Intestinal fungal burden represented as CFU-g^-1^ stool. (E) Representative flow cytometry plots examining IgA bound *C. glabrata* directly ex vivo from cecal contents quantified in (F) as frequency IgA bound of anti-*Candida*^+^ cells. (G) IgA Geomean of day 28 cleared intestinal content wash from mono-colonized gnotobiotic mice incubated with indicated cultured strains. (B) Ordinary one-way ANOVA with Dunnett’s multiple comparisons test, n=4-9, comparing Cg27 to all other strains. Data are mean +/- SEM. (D, F, G) Two-way ANOVA with Tukey’s multiple comparison test. n=5-9 mice per group. Each dot represents an individual mouse across three independent experiments.

At day 28, we assessed IgA reactivity toward individual strains within each pool. Intestinal IgA bound 22 strains at levels comparable to or exceeding Cg1, while 11 strains, including all pool 1 strains, showed no detectable IgA binding (Fig. 1B). Thus, the ability to stimulate IgA varies substantially among *C. glabrata* strains. Cg27 was the most strongly IgA-targeted strain, exhibiting 2–3-fold higher IgA binding than Cg1 and other strains. Cg27 is also part of pool 4, which elicited the most rapid IgA response (Fig. S1A). We next tested whether Cg27, like Cg1, could independently induce IgA in monocolonized germ-free (GF) mice (Fig. 1C). We compared this response to CBS138, the genome reference strain, which we found was a poor IgA inducer in preliminary studies. Cg1, Cg27, and CBS138 achieved similar gut burdens, although Cg1 and Cg27 trended toward higher small intestinal colonization than CBS138 (Fig. 1D). Strikingly, nearly all intestinal Cg1 and Cg27 cells were IgA-bound, compared with only ∼30% of CBS138 cells (Fig. 1E-F).

Free intestinal IgA from Cg1- and Cg27-colonized mice was highly reactive to cultured Cg1 and Cg27 and showed strong cross-reactivity between these strains, but significantly less binding to CBS138. In contrast, IgA from CBS138-colonized mice did not react against any of these strains (Fig. 1G, Fig. S1B). Intestinal IgG1 responses were detectable but substantially weaker than IgA and preferentially targeted Cg27 (Fig. S1C). Serum analyses mirrored mucosal findings, with Cg1 and Cg27 inducing strong strain-reactive and cross-reactive IgA and IgG responses, again most reactive against Cg27 (Fig. S1D). CBS138 did not induce a *C. glabrata-*reactive serum response.

We also investigated whether strain-dependent IgA induction was conserved across mouse genetic backgrounds. Like we observed in B6 mice, Cg27, but not CBS138, induced a Cg27-specific mucosal IgA response in antibiotic-treated Swiss Webster (SW) mice (Fig. S2A-E). Collectively, these data demonstrate that both the induction and specificity of *C. glabrata*-induced IgA are highly strain dependent. This also identifies Cg1 and Cg27 as immunogenic strains that drive robust, cross-reactive antibody responses in the gut, and CBS138 as a poor IgA inducer.

### Mucosal IgA targets putative adhesin Awp11

To investigate how IgA-reactive strains differ from non-reactive strains, we performed whole-genome sequencing on CBS138 and clinical isolates in our collection (Cg7 was excluded due to failed library preparation). Phylogenetic analysis showed that strains do not cluster by IgA reactivity (Fig. 2A), as measured by our pooled IgA screen in Fig. 1A, indicating that mucosal IgA does not preferentially target specific genetic clades of *C. glabrata*. To assess whether IgA-inducing and non-inducing strains differ transcriptionally, we performed RNA-seq on two IgA-reactive strains (Cg1, Cg27) and three non-reactive strains (CBS138, Cg4, Cg20), with the latter two selected for genetic similarity to the most highly IgA-targeted strain, Cg27 (Fig. 2A). Pairwise comparisons identified numerous differentially expressed genes between strains, with fewest differentially expressed genes observed when comparing strains with highest genetic similarity (e.g. Cg27 and Cg20) (Fig. S3A-B, Table. S2). However, when comparing all IgA-reactive versus non-reactive strains, a single gene, *AWP11* (CAGL0J12067), notably upregulated, with up to 50-fold higher expression, in IgA-inducing strains (Fig. 2B).

**Figure 2.**
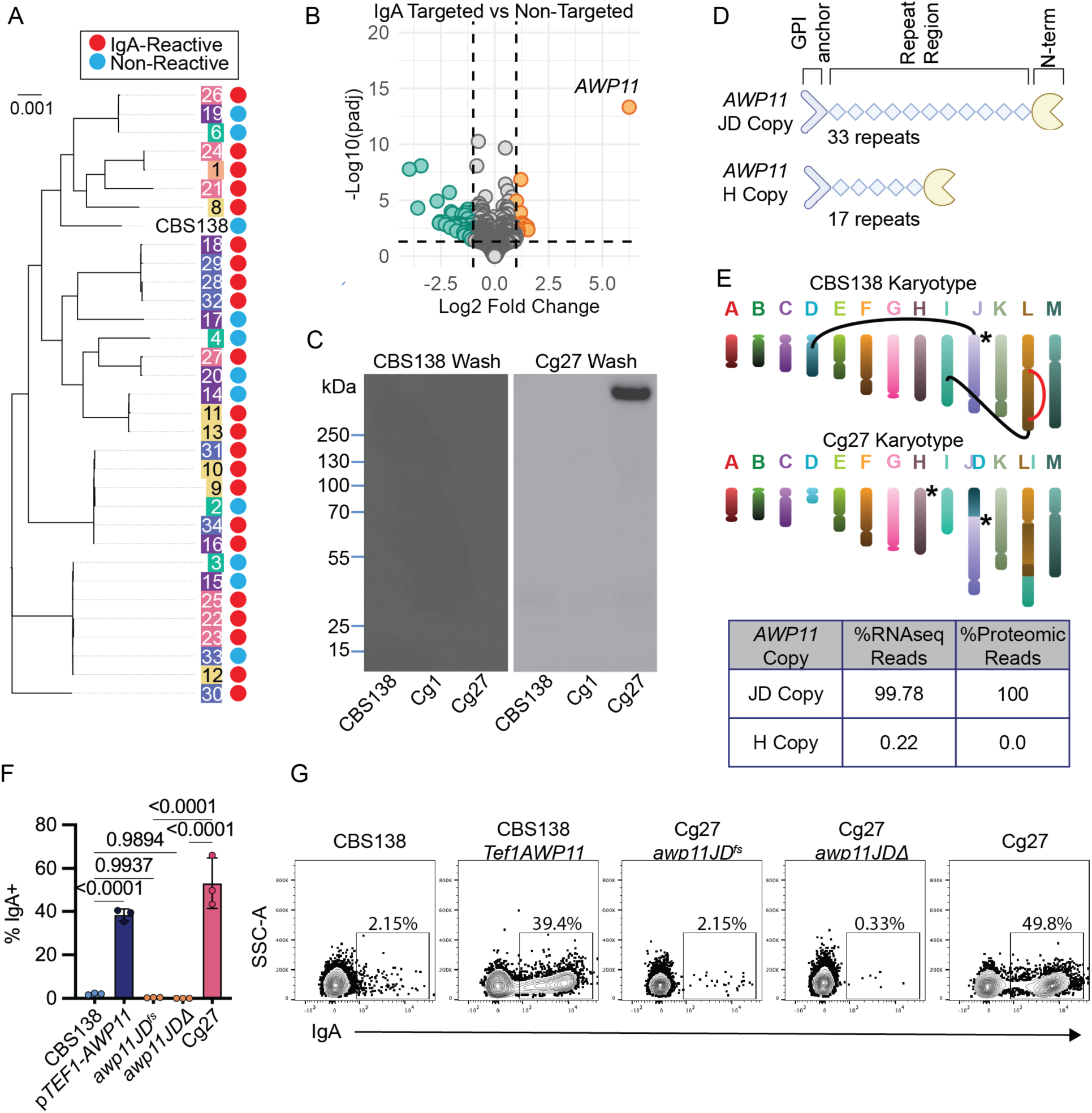
Mucosal IgA targets putative adhesin Awp11. (A) *C. glabrata* clinical isolate phylogeny denoted by pooled screen colors and IgA-reactivity status. (B) In vitro *C. glabrata* RNA-seq Volcano plot comparing IgA-targeted and non-targeted strains. (C) CBS138, Cg1, and Cg27 cell wall Western blotted with cleared intestinal wash from mice monocolonized with CBS138 or Cg27 (D) Schematic demonstrating the predicted protein structure of Awp11 in all *C. glabrata* isolates and the differences between copies within Cg27. (E) Schematic of the chromosomal structure of CBS138 and Cg27. Black lines indicate translocations, red lines indicate inversions, and * indicate the location of *AWP11*. Proportion of sequenced RNA reads from (B) and peptide fragments from (C) that aligned exclusively to each copy of *AWP11* in Cg27. (F) Frequency and (G) representative flow plots of IgA binding to indicated strains following incubation with cleared intestinal wash from mice monocolonized with Cg27/ (F) Two-way ANOVA using Tukey’s multiple comparisons test. n=3.

In parallel, we sought to identify IgA-targeted protein antigens expressed on the cell surface of Cg1 and Cg27. We isolated cell wall-associated proteins from Cg1, Cg27, and CBS138 and performed a western blot using cleared intestinal wash from GF mice or mice monocolonized with each strain. As expected, IgA from GF or CBS138-colonized mice showed low reactivity against *C. glabrata* cell wall proteins (Fig. S3C-F, Fig. 2C). IgA induced by Cg1 recognized a broad range of proteins (∼275–300 kDa) without clear enrichment relative to CBS138 (Fig. S3E). In contrast, Cg27-induced IgA specifically recognized a single ∼300 kDa protein present only in the Cg27 cell wall (Fig. 2C, Fig. S3F). LC–MS/MS analysis of proteins in this size range identified the putative adhesin Awp11 as the only protein unique to Cg27 (Table. S3). *AWP11* encodes a predicted GPI-anchored adhesin belonging to the recently described cluster V family (14), characterized by an N-terminal ligand-binding domain, a serine/threonine-rich repeat region, and a C-terminal GPI anchor (Fig. 2D). While related adhesins (e.g., Awp2) have been implicated in adherence to abiotic surfaces (14), the function of Awp11 remains uncharacterized.

Adhesin gene content and copy number has been shown to vary between *C. glabrata* strains (15), so we investigated whether *AWP11* copy number is associated with expression levels. All strains in our collection encode at least one copy of *AWP11* with 13/33 having two copies. Phylogenetic relatedness correlated closely with *AWP11* copy number. Cg27 (high *AWP11* expression) and Cg20 (low *AWP11* expression) have two copies, while CBS138 (low *AWP11* expression) and Cg1 (high *AWP11* expression) have only one (Fig. S4A-C). This indicates that while *AWP11* copy number varies between strains, it is unlikely to be responsible for strain-dependent differences in *AWP11* transcription.

The majority of *C. glabrata* adhesin genes are subtelomeric and subjected to epigenetic silencing, which has been associated strain-dependent variability in adhesin expression (16, 17). Therefore, we asked whether *AWP11* genomic location might explain differential expression. Notably, up to 34% of the top 50 differentially expressed genes between strains encode predicted adhesins (Fig. S3G), including *AWP11*, which is located in CBS138 on the right subtelomere of chromosome J. To assess structural variations in these regions, we generated de novo genome assemblies for CBS138, Cg1, and Cg27 (Fig. S5A-D). Cg1 genome structure was similar to CBS138, both encoding a single *AWP11* copy at the subtelomere of chromosome J (Fig. S5C). In contrast, Cg27 exhibited three major chromosomal rearrangements compared to CBS138, including a fusion between the left arm of chromosome D and the right arm of chromosome J. This J/D fusion is directly adjacent to one *AWP11* copy in Cg27. Cg27’s second *AWP11* copy resides at the subtelomere of chromosome H (Fig. 2E, Fig. S5E).

We hypothesized that this J/D chromosomal rearrangement in Cg27 has resulted in the de-repression the *AWP11* gene copy adjacent to this fusion. Sequence polymorphisms between the two Cg27 alleles allowed us to assign transcripts and peptides to each copy. 99.78% of *AWP11* transcripts and 100% of detected Awp11 peptides mapped to the allele adjacent to the J/D fusion (Fig. 2E), indicating that nearly all *AWP11* transcripts and protein is expressed from this copy. Cg20, the closest genetic relative to Cg27, also encodes two *AWP11* copies but is not targeted by intestinal IgA, and expresses very low levels of *AWP11* in culture (Fig. 1B, Fig. S3A). Alignment of Illumina reads confirmed that Cg20 lacks reads spanning this J/D fusion, whereas these can be found in Cg27 (Fig. S5E). These data support that this structural rearrangement has resulted in constitutive *AWP11* expression in Cg27.

To test whether Awp11 is the dominant IgA target, we generated two loss-of-function mutations specifically in the J/D-associated *AWP11* allele in Cg27 (Fig. 2F-G): a frameshift insertion introducing a premature stop codon (*awp11JD^fs^)* using CRISPR (18), and a disruption of the first 496bp of the gene (*awp11JDΔ*). Both mutations abolished IgA binding to Cg27, demonstrating that Awp11 is required for IgA recognition. We were unable to disrupt *AWP11* in Cg1 despite multiple attempts. Despite this, we found that Cg1-induced IgA failed to recognize the Cg27 *awp11JD^fs^*, indicating that Cg1 also elicits Awp11-reactive IgA (Fig. S3H). However, differences in western blot patterns suggest that Cg1- and Cg27-induced IgA recognize distinct Awp11 epitopes masked by the linear versus 3D nature of western and flow epitopes respectively, or that Cg1-induced IgA recognizes other cell wall proteins in addition to Awp11 (Fig. S3F-G).

Finally, we asked whether inducing *AWP11* transcription in CBS138 is sufficient to drive IgA binding when incubated with Cg27-induced IgA. We replaced the native *AWP11* promoter in CBS138 with the constitutive pTEF1 promoter. This resulted in resulted in strong binding by Cg27-induced IgA, whereas parental CBS138 remained non-reactive (Fig. 2F-G). Together, these data identify Awp11 as a dominant target of intestinal IgA and demonstrate that in Cg27, a chromosomal rearrangement is likely responsible for increased *AWP11* expression.

### Awp11 is necessary for adaptive immune responses in vivo

To determine whether Awp11 is necessary for IgA induction in vivo, we colonized antibiotic-treated mice with CBS138, Cg27, or Cg27 *awp11JD^fs^* and measured the dynamics of IgA targeting of fecal *C. glabrata* directly ex vivo, as well as levels of free *C. glabrata-*reactive IgA. We found that ∼20-30% of all strains were IgA bound in vivo within 6 hours of colonization within the feces, and the percentage of IgA binding decreased over the first 7 days (Fig. 3A). This early IgA targeting occurred despite undetectable *C. glabrata*–specific IgA (free IgA reactive against *C. glabrata* from culture) prior to inoculation (Fig. 3B), suggesting that this early targeting may be non-specific. Consistent with this, staining for major cell wall components (β-glucan, chitin, and mannan) revealed dynamic changes in surface exposure during the first 48 hours, as previously described (19), which stabilized by days 5–7 (Fig. S6A–C). This stabilization coincided with reduced IgA binding, supporting the idea that transient cell wall remodeling may lead to non-specific recognition by endogenous IgA.

**Figure 3.**
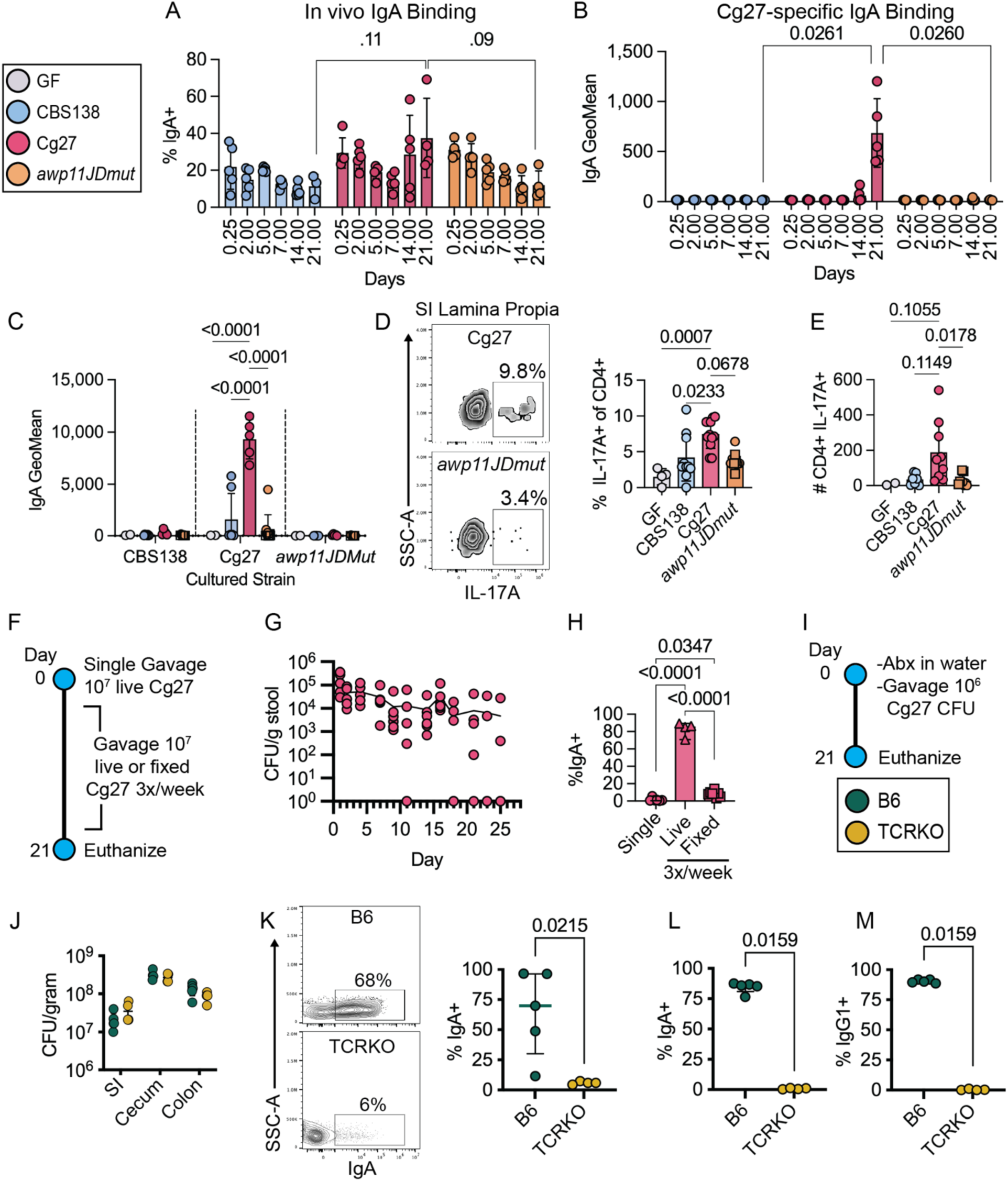
Mucosal adaptive immune responses are Awp11 dependent. (A) Time course of in vivo *C. glabrata* IgA binding frequency in stool and (B) IgA geomean of cultured Cg27 incubated with paired stool wash from antibiotic treated mice. n=5 mice per group. (C) IgA Geomean of day 21 SI content wash from mono-colonized gnotobiotic mice incubated with indicated cultured strains. (D) Representative flow cytometry plots and frequency of IL-17A+ cells as a percentage of CD4+ T cells in Cg27 (top) and *awp11JDmut* (bottom) gnotobiotic mouse SI lamina propria. (E) Total cell number of CD4+ IL-17A+ in SI lamina propria. n=4-13 mice per group, from 4 independent experiments. *awp11JDmut* includes both *awp11JD^fs^* (circles), *awp11JD*Δ (squares). (F) Timeline for non-antibiotic single gavage vs repeated live or formalin-fixed Cg27 gavage experiment. (G) Cg27 stool fungal burden over time in non-antibiotic treated mice gavaged with 10^7^ CFU. (H) Day 21 SI wash IgA binding frequency to in vitro cultured Cg27. 4-8 mice per group from one independent experiment. (I) Schematic for TCRKO antibiotic treated colonization experiment. (J) Cg27 fungal burden in intestinal contents. (K) Representative flow plots and quantification of IgA binding in SI contents in vivo at day 21. Frequency of day 21 (L) SI content wash IgA and (M) serum IgG1 binding to in vitro grown Cg27. n=5 mice per group. All data are mean +/- SD. Each data point represents an individual mouse. (A-C, J) Two-way ANOVA, (D, E) Kruskal-Wallis with Dunn’s Multiple Comparison test, (H) One-way ANOVA with Tukey’s multiple comparison test, (K-L) Two-tailed Mann-Whitney test.

Beginning at day 14, IgA binding increased specifically for Cg27, but not for CBS138 or Cg27 *awp11JD^fs^*, and was also elevated in small intestinal contents at day 21 (Fig. S6E). This increase coincided with a robust rise in free Cg27-specific IgA (Fig. 3B). Notably, the Cg27 *awp11JD^fs^* failed to induce IgA responses against either itself or Cg27 (Fig. S6D), demonstrating that Awp11 is required for IgA induction. However, constitutive expression of *AWP11* in CBS138 (p*TEF1-AWP11*) did not induce an IgA responses under antibiotic-treated conditions (Fig. S6F), indicating that Awp11 expression may not be sufficient for CBS138 to induce an IgA response. Similar results were observed in germ-free mice, where Cg27, but not the *awp11JD^fs^* or *awp11JDΔ,* induced a specific IgA response despite comparable total IgA levels across groups (Fig. 3C, Fig. S6G–I). Together, these data show that Awp11 is necessary for induction of Cg27-specific IgA, but may not be sufficient in other *C. glabrata* strains.

We next examined mucosal T and B cell responses in germ-free mice monocolonized with each strain, focusing on key sites of IgA-induction and IgA-production including the small intestinal lamina propria (LP), Peyer’s patches (PP), and mesenteric lymph nodes (MLN) (20) (Fig. S7A). Cg27 colonization led to increased frequencies of IL-17A producing CD4+ T cells in the SI LP compared to CBS138 and the *awp11JDmut* (Fig. 3D–E), with similar trends observed in PP and MLN (Fig. S7B–E). Frequencies of RORγt+ CD4+ T cells were also elevated in Cg27-colonized mice although all *C. glabrata* strains increased RORγt+ cells relative to germ-free controls (Fig. S7F). Notably, despite robust IgA induction in Cg27 mice, we found no differences in germinal center B cell cells (Fig. S7G-H) or CXCR5+ PD1+ (T follicular helper) cells (Fis. S7I) in gut associated lymphoid tissues. Additionally, we report no differences in the percentages of FOXP3+ (T regulatory) or IFNγ+ (Th1) in the gut associated lymphoid tissues (GALT) or SI LP in all groups (Fig. S7J-K). This indicates that Awp11 is necessary for both mucosal antibody and Th17 cell expansion in the SI.

Because antibiotic treatment can disrupt the mucosal barrier (21), we next tested whether Cg27 induces IgA responses in the absence of antibiotics and whether viability is required. SPF mice were gavaged either once or repeatedly (3x/week for 21 days) with live or formalin-fixed Cg27 (Fig. 3F). Importantly, fixed Cg27 retained IgA binding by Cg27-induced IgA, indicating that fixation does broadly alter the antigenicity of Awp11 (Fig. S6J). A single gavage resulted in persistent colonization (Fig. 3G) but did not induce a detectable IgA response. In contrast, repeated gavage with live Cg27 elicited robust Cg27-specific IgA, whereas fixed cells failed to do so (Fig. 3H), indicating that sustained exposure to viable fungi is required.

Microbiota-targeting IgA can be induced through T cell-dependent and independent pathways (20). To determine whether IgA induction is T cell–dependent, we compared responses in antibiotic-treated C57B6/J and *Tcrb^-/-^d^-/-^* (TCRKO) mice colonized with Cg27 Fig. 3I). TCRKO IgA plasma cells undergo no somatic hypermutation or affinity maturation in the gut but are still capable of IgA class switching (13, 22, 23). Cg27 colonized both groups similarly (Fig. 3J). Cg27-specific mucosal IgA, systemic IgG, and IgA coating of fungi were observed in WT mice (Fig. 3K–M) but were absent in TCRKO mice. This demonstrates that T cells are required for Cg27-specific antibody induction.

### Cg27-induced antibody responses do not protect from systemic infection

Commensal *C. albicans* induces antibody and T cell responses provide protection against subsequent disseminated infection (24–28). Therefore, we tested whether colonization-induced immunity protects against systemic *C. glabrata* infection. Unlike *C. albicans,* systemic challenge with *C. glabrata* is non-lethal in mice (29), so we used fungal burden as a proxy for virulence/fungal fitness in this model. Wild-type and μMT^-/-^ mice, which lack mature B cells and fail to mount an intestinal IgA response against Cg27 (Fig. S10H) were colonized with CBS138 or Cg27 for 21 days and then challenged intravenously with each strain (Fig. S8A). Despite robust antibody responses in Cg27-colonized wild-type mice, fungal burdens in spleen, liver, and kidney were comparable between μMT^-/-^ and WT mice (Fig. S8B). Although potentially impaired (30), μMT^-/-^ mice can develop antigen-specific T cell responses which may explain equal fungal clearance. To test this, we colonized WT mice with either CBS138 or Cg27 for 21 days, or left them uncolonized, and then intravenously challenged all mice, reasoning that those left uncolonized with be immunologically naïve to *C. glabrata* prior to disseminated infection. Again, we found no difference in fungal burden in pre-colonized vs uncolonized mice (Fig. S8C). This suggests that mature T and B cell responses, including the Awp11-specific IgA response, may be ineffective in limiting a systemic infection with these strains.

### Awp11 does not enhance *C. glabrata* adherence to intestinal epithelial cells

Commensal bacteria that closely associate with the intestinal epithelium have been shown to be strong inducers of IgA and Th17 responses (12, 13). To test whether Awp11 immunogenicity reflects enhanced ability for Cg27 to adhere to the intestinal epithelium, we quantified tissue-associated fungal burdens in antibiotic-treated mice colonized with CBS138, Cg27, or the Cg27 *awp11JD^fs^.* We focused on early time points prior to the development of anti-Cg27 IgA to avoid the potential effect of Cg27 and Awp11-specific antibodies altering adherence. At day 8 post-colonization, Cg27 reached ∼10 fold higher burdens than CBS138 in the small intestine lumen and tissue (though statistically trending at the tissue), and a ∼2 fold higher burden in the colon contents. Colonization of the Cg27 *awp11JD^fs^*mutant in the SI and colon lumen was statistically lower than Cg27, though only modestly, and there was no significant difference in tissue-associated burdens with this mutant (Fig. S9A). To directly assess the role of IgA, we compared colonization in antibiotics-treated WT and µMT^⁻/⁻^ mice. After 21 day, tissue-associated burdens of Cg27 were unchanged between WT and µMT^⁻/^, indicating that IgA does not restrict epithelial association under these conditions (Fig. S9B). Together, Awp11 is not required for close colonization of the intestinal tissue in vivo.

We next tested whether Awp11 promotes epithelial adhesion in vitro using Caco-2 human colon epithelial cells. Cg27 exhibited a trending, but not significant, increase in adherence relative to CBS138. However, Cg27 *awp11JD^fs^* showed significantly greater adherence to Caco-2 cells than both Cg27 and CBS138 (Fig. S9C), suggesting that Awp11 may limit, rather than promote, adhesion to Caco-2 cells. Addition of IgA purified from GF or Cg27-colonized mice did not alter adhesion (Fig. S9C). Together, these data indicate that Awp11 may simply function as an intestinal epithelial cell targeting adhesin, and the intestinal IgA does not impact *C. glabrata* adherence to intestinal epithelial cells.

### Awp11 confers competitive advantage during gut colonization

Fungal effectors, including *C. albicans* hyphal formation and candidalysin, are dispensable for colonization of antibiotics-treated or GF, but be critical for fitness in a complex microbiota (31). Therefore, we hypothesized that the function of Awp11 may depend on the gut microbial community. To test this, we compared colonization of the Cg27 and *awp11JD^fs^*in SPF mice, finding that the mutant colonized mice to a significantly lower burden that Cg27, and was cleared by day 15 (Fig. S9D). To more directly assess fitness, we performed competitive colonization experiments between Cg27 and the *awp11JD^fs^* or the *awp11JDΔ* mutants. In antibiotic-treated mice, Cg27 displayed a modest but significant fitness advantage over both *awp11JD^mut^*strains (data combined for both competitions in Fig. 4A). After 21 days, this advantage was evident across most intestinal sites, though less pronounced in small intestinal tissue (Fig. 4D). In contrast, in SPF mice, Cg27 rapidly outcompeted both Cg27 *awp11JD* mutant strains within three days (Fig. 4B) and remained the dominant strain in all intestinal regions by day 21 (Fig. 4E). We also performed this competition in GF mice without antibiotic-treatment, finding no competitive difference between strains (Fig. 4C). Together, these findings demonstrate that Awp11 enhances *C. glabrata* fitness and persistence specifically in the context of a complex microbial community.

**Figure 4.**
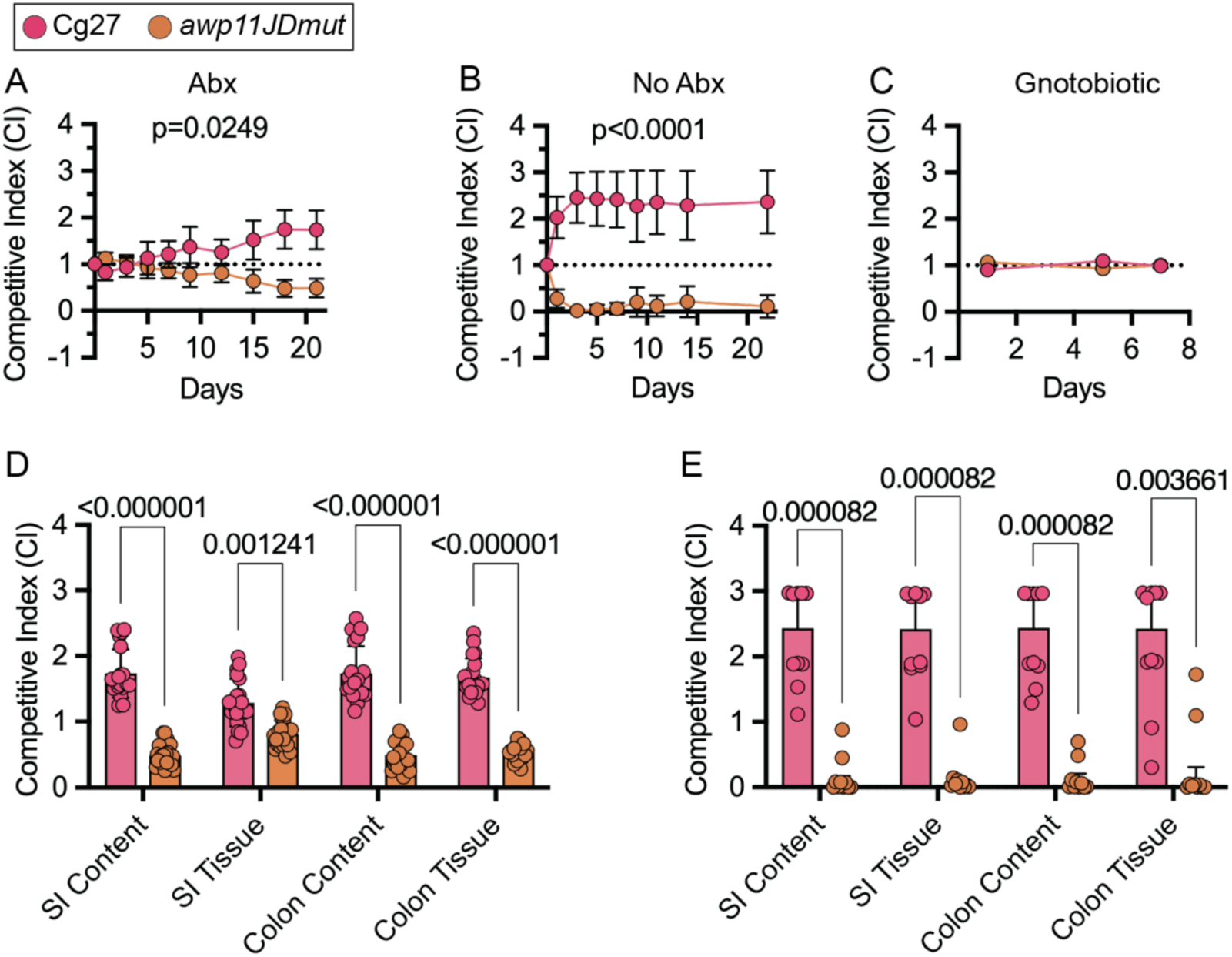
Awp11 confers competitive advantage during gut colonization. Competitive index (CI) of Cg27 and Cg27 *awp11JD^fs^* in WT SPF mice treated with (A) or without (B) antibiotic treatment. (C) CI of Cg27 and Cg27 *awp11JD^fs^* in GF mice. Day 21 CI of Cg27 versus Cg27 *awp11JD^fs^* in intestinal contents and tissues in antibiotic treated (D) and non-antibiotic treated mice (E). (A-E) Paired *t*-test. n=10 mice per group. Data are mean +/- SD.

### IgA alters the competitive fitness of Awp11-expressing *C. glabrata*

To determine whether IgA influences Cg27 Awp11-dependent colonization, we compared the competitive fitness of Cg27 and Cg27 *awp11JD^fs^*in WT and *Igha^-/-^* (IgA^-/-^) mice. Because Cg27 elicits a specific IgA response only in GF or antibiotic-treated animals (Fig. 3H), we started the competition under antibiotic treatment (Fig. 5A). B6, but not IgA^-/-^ mice developed a robust Cg27-reactive IgA response which peaked at day 25 (Fig. 5B-C). We did not observe a compensating IgG or IgM response in IgA^-/-^ mice by day 25 (Fig. 5G and data not shown). Like what we observed in Fig. 4A, Cg27 has a modest competitive advantage in WT but not IgA^-/-^ mice during antibiotic treatment, and this advantage was not altered after the development of Cg27-specific IgA in WT mice (Fig. 5B-C). We verified that this was not due to the selection of de novo Cg27 mutants that avoid IgA detection, finding that Cg27 fecal isolates retained IgA reactivity, while *awp11JD^fs^* isolates remained non-reactive at day 25 (Fig. 5D).

**Figure 5.**
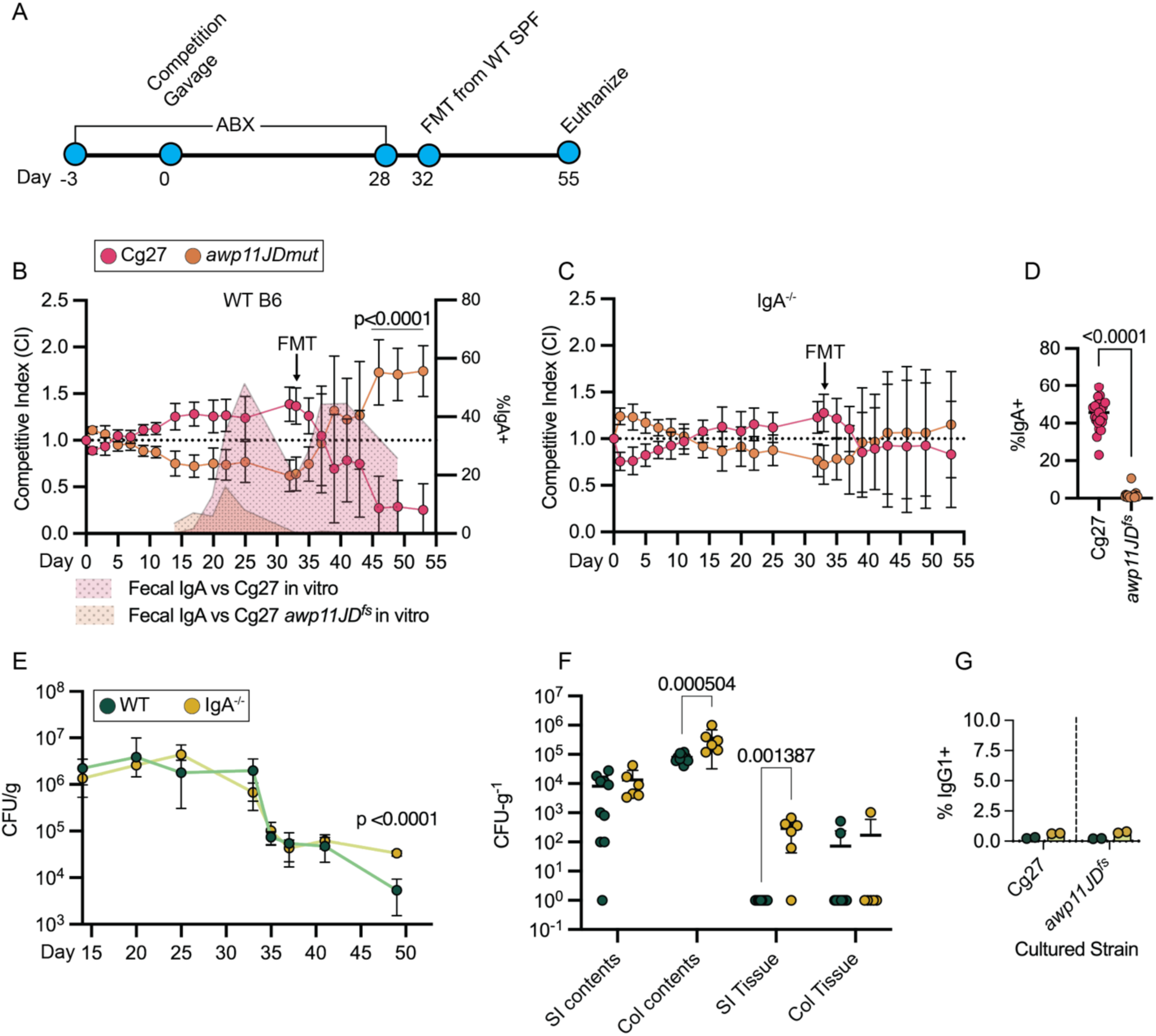
IgA alters the competitive fitness of Awp11-expressing *C. glabrata*. (A) Schematic of antibiotic-FMT competition experimental design. (B) CI of Cg27 and Cg27 *awp11JD^fs^* in WT mice. Levels of Cg27 and Cg27*awp11JD^fs^* -reactive IgA in cleared fecal wash over time indicated in shaded overlay. (C) CI of Cg27 and Cg27 *awp11JD^fs^* in IgA^-/-^ mice. n=6-10 mice per group. (D) IgA reactivity to indicated gut-passaged strains from culture after incubation with cleared intestinal wash from mice monoclolonized with Cg27. n=23 of each strain (E) Fecal fungal burden (F) Intestinal content and tissue fungal burden at 55 days post colonization. n=6-10 mice per group. Data are mean +/- SD. (G) Intestinal IgG1 reactivity against Cg27 and Cg27 *awp11JD^fs^* from culture. Cleared fecal wash from WT and IgA^-/-^ mice at day 25 post colonization. n=2 (B) *Paired t-* test. Before FMT, p = 0.0059, and after FMT, p = 0.1036. Data are mean +/- SD. Additional p values shown over days 46, 49, and 53 were calculated by ordinary two-way ANOVA using Šídák’s multiple comparisons test. (C) *Paired t-* test. Before FMT, p = 0.9049, and after FMT, p = 0.6770. Data are mean +/- SD. (D) Mann-Whitney U test. (E) Mixed-effects model with Geisser-Greenhouse correction and Tukey’s multiple comparisons test. (F) Mann-Whitney U test. Each dot represents an individual mouse.

To assess whether restoring the microbiota alters the competitive fitness of Cg27 and *awp11JD^fs^,* antibiotics were withdrawn at day 28 and mice received a fecal microbiota transfer (FMT) on day 32, from SPF B6 donors (Fig. 5A). Following microbiota reconstitution, Cg27 *awp11JD^fs^* outcompetes Cg27 in WT mice by day 55, but not in IgA^-/-^ mice (Fig. 5B-C). Notably, restoration of the microbiota in IgA^-/-^ mice did not result in clearance of *awp11JD^fs^* like we observed in naïve SPF animals (Fig. 4B), suggesting either key bacterial species that drive Awp11-dependent colonization of SFP mice are not fully restored post FMT, or that Awp11 may be functionally important in establishing colonization, but less important for persisting once established.

Finally, Following FMT, total fungal burden decreased by ∼100-fold in all mice (Fig. 5E). However, IgA^-/-^ mice maintained significantly higher *C. glabrata* burdens (including both strains) at endpoint in stool, colon contents, and small intestinal tissue (Fig. 5E–F). Notably, *C. glabrata* was undetectable in the small intestinal tissue of 8/9 WT mice, whereas all IgA^-/-^ mice retained detectable colonization at this site (Fig. 5F). Collectively, this suggests that the microbiota synergizes with IgA to limit the fitness of Awp11-expressing *C. glabrata*.

### Intestinal environment promotes *AWP11* expression in clinical isolates

High *AWP11* expression in Cg27 is likely the result of the chromosome J-D fusion event that is specific to this strain in our collection. However, *AWP11* genes are typically subtelomeric in most *C. glabrata* strains and subjected to epigenetic silencing mediated by the Sir complex and other chromatin regulators (32, 33). We tested whether Sir-mediated repression occurs in CBS138 using a *sir3Δ* mutant. Loss of *SIR3* increased *AWP11* transcript levels ∼1000 fold and altered expression of several other cluster V adhesins (Fig. 6A, Fig. S10A-D). The *sir3Δ* mutant also showed increased IgA targeting by Cg27-induced IgA, though less than Cg27, suggesting additional regulatory mechanisms affecting Awp11 expression or localization in CBS138 (Fig. 6B). In contrast, *sir3Δ* in Cg27 did not significantly affect *AWP11* transcript levels or IgA binding, consistent with the J/D copy being uncoupled from this subtelomeric regulation (Fig. 6A-B, Fig S5).

**Figure 6.**
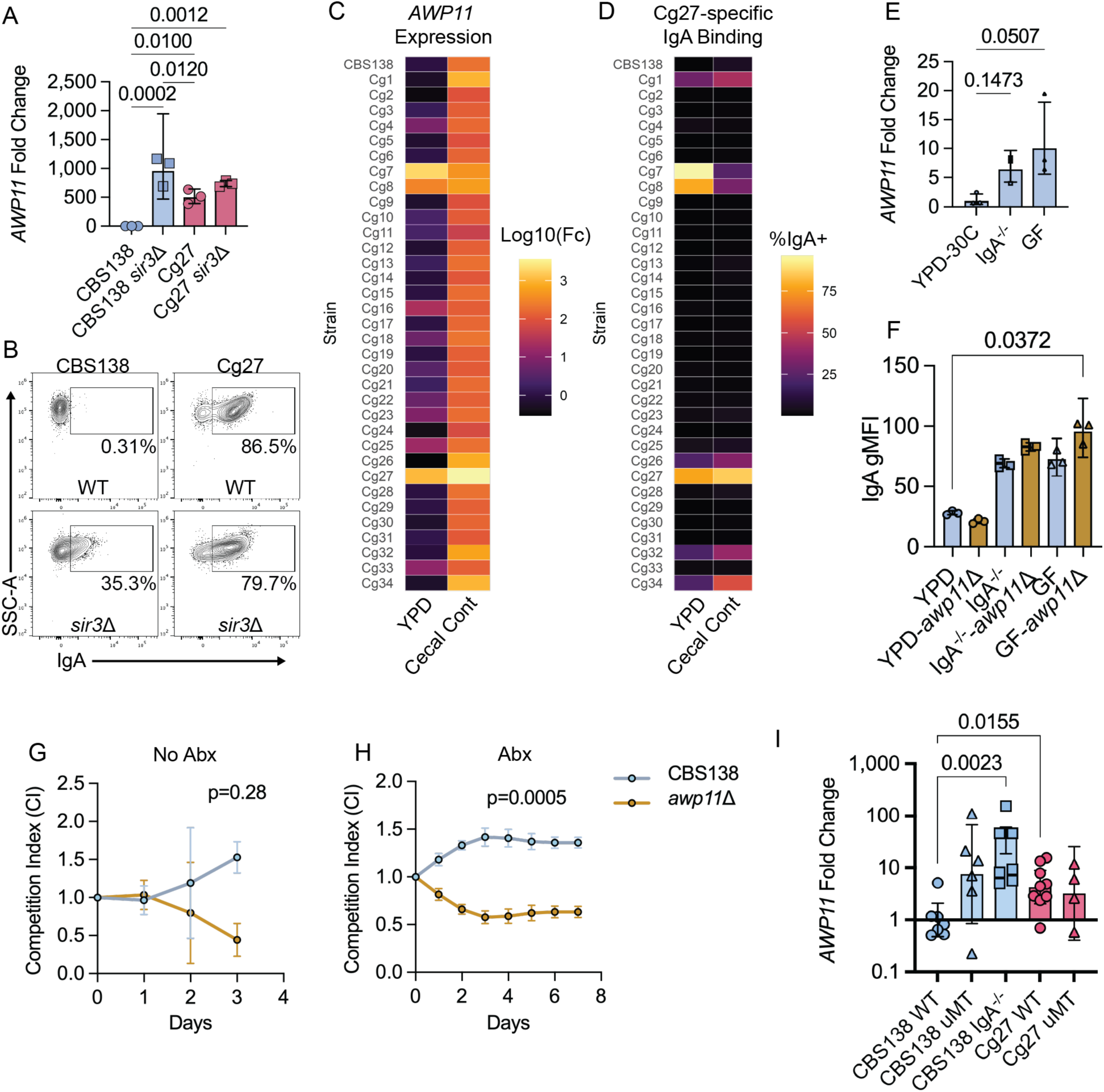
Intestinal environment promotes *AWP11* expression. (A) *AWP11* qRT-PCR analysis of WT and *sir3Δ* CBS138 and Cg27 grown in vitro at 30°C in YPD. (B) Representative flow cytometry plots of IgA binding to WT and *sir3Δ* CBS138 and Cg27 grown in vitro following incubation with Cg27 monocolonized GF SI contents. (C-D) *C. glabrata* clinical isolates grown for 3 hours in YPD at 30°C or IgA^-/-^ or GF mouse cecal contents at 37°C. (C) Heatmap of *AWP11* expression measured by qRT-PCR. Data are represented as fold change relative to an internal housekeeping gene, *ACT1*, and CBS138 grown in YPD sample. (D) Heatmap of IgA binding frequency following incubation with cecal wash supernatant from gnotobiotic mice colonized with Cg27 for 21 days. (E) CBS138 *AWP11* expression measured by qRT-PCR following incubation in YPD or IgA^-/-^ or GF cecal contents. (F) IgA geomean of CBS138 and CBS138 *awp11 Δ* strains following incubation with cecal wash supernatant from gnotobiotic mice colonized with Cg27 for 21 days. n=3. Competition between CBS138-GFP or CBS138-iRFP and CBS138 *awp11 Δ* in (G) non-antibiotic treated and (H) antibiotic treated B6 mice. n=10 mice per group from two independent experiments. (I) Day 21 *AWP11* expression, measured from cecal content *C. glabrata,* represented as fold change relative to CBS138 WT group. n= 5-9 per group. Data are mean +/- SD. Each data point represents an individual mouse. (E) Kruskal-Wallis with Dunn’s Multiple Comparison, (F) One-way ANOVA with Tukey’s multiple comparison test, (G-H) Paired two-tailed t-test, (I) Welch’s ANOVA with Holm-Sidáks multiple comparisons test.

Sub telomeric adhesin genes can be unsilenced in vivo during growth in various in vivo-mimicking conditions (34). We investigated whether in non-Cg27 strains, *AWP11* expression could be turned on in response to signals present in the intestinal tract. We compared *AWP11* transcripts by qPCR after culture in standard YPD media at 30°C or cecal contents from IgA^-/-^mice at 37 °C. This method has been utilized before to mimic growth in intestinal conditions and was used to induce hyphae formation in *C. albicans* with similar frequencies to intestinal colonization (31). Use of the IgA^-/-^ contents allows us to analyze Awp11 cell wall expression by probing with Cg27-induced intestinal IgA. In standard YPD media, 76% of strains expressed low levels of *AWP11* like the CBS138 strain, but 3 strains, including Cg27, had over 100-fold increase. Strikingly, all strains grown in cecal contents had a >100 fold increase in *AWP11* expression relative to CBS138 in YPD (Fig. 6C). Additionally, all but 2 strains displayed increased *AWP11* expression in cecal contents relative to their YPD cultures (Fig. S10E). We also observed increased binding by Cg27-specific IgA in most strains, though notably, this increase in IgA recognition was more modest than expected from the transcriptional induction (Fig. 6C, Fig S10F).

We investigated whether microbial signals and IgA influence *AWP11* expression in CBS138. In this strain, we found that relative to YPD, cecal contents from GF and IgA^-/-^ mice induced *AWP11* transcription similar to contents from SPF mice, indicating that IgA and the microbiota are not necessary for inducing transcription (Fig. 6E). We also found increased Cg27-induced IgA reactivity to CBS138 when grown in all cecal contents compared to YPD (Fig. 6F). To investigate whether the increase in IgA binding to CBS138 was Awp11-specific, we repeated this experiment with a CBS138-*awp11Δ.* Interestingly, there were no differences between CBS138 and CBS138-*awp11*Δ in IgA recognition, suggesting that there may be upregulation of other antigens that cross react with Cg27-induced IgA (Fig. 6F). To support this, we analyzed CBS138 expression of other *AWP* genes finding increased expression of *AWP2, AWP4, and AWP10* in GF contents relative to YPD (Fig. S10G-J).

These data show that CBS138 may turn on *AWP11* expression in response to intestinal cues, and therefore, may be functionally important in the gut. To test this, CBS138 and CBS138-*awp11*Δ were competed in both SPF and antibiotic-treated mice. Notably, both strains were undetectable by day 4 in SPF mice, limiting our ability to fully assess the function of CBS138 Awp11 in colonization of a complex microbiota. Despite this, we found that the CBS138 *awp11*Δ was significantly less fit than CBS138 in both groups of mice (Fig. 6 G-H). These data show that CBS138 turns on *AWP11* expression in response to intestinal conditions in vitro and requires *AWP11* for competitive fitness in vivo. However, CBS138 generally fails to induce Awp11-reactive IgA, with notable but infrequent responses detected in 3/16 GF mice (Fig. 1G, Fig. 2C), suggesting that *AWP11* is not constitutively expressed in vivo.

We next asked whether IgA influences CBS138 and Cg27 *AWP11* expression in vivo. *AWP11* expression in cecal content was analyzed from B6, IgA^-/-^, and μMT^-/-^ mice colonized with CBS138 or Cg27 for 21 days. No difference was observed in fecal fungal burden within fungal strains and mouse backgrounds (Fig. S10K). As expected, Cg27 was more highly bound by IgA in the gut compared to CBS138 in B6 mice, and no IgA binding to Cg27 was found in μMT^-/-^ mice (Fig. S10I). Strikingly, CBS138 *AWP11* expression was up to 100-fold higher in μMT^-/-^ and IgA^-/-^ mice relative this strain in B6. This suggests that CBS138 *AWP11* expression is suppressed in the presence of IgA (Fig. 6I). Cg27 *AWP11* expression was similarly high in both WT and μMT^-/-^ mice, consistent with its genetic chromosomal fusion event resulting in constitutive *AWP11* expression. These results suggest that *C. glabrata* dynamically regulates *AWP11* expression to balance intestinal fitness in a complex microbiota and immune evasion in the gut.

## Discussion

Using a collection of bloodstream isolates we identified an adhesin-like protein, Awp11, that is specifically targeted by T cell-dependent IgA antibody responses throughout the intestinal tract.

Awp11 expression drives strain-to-strain variability in IgA responses and provides a fitness advantage in a complex microbiota. In the majority of tested strains, *AWP11* expression is induced within the intestinal environment, though it may be specifically regulated by IgA in the gut. Together, these findings identify Awp11 as both a key intestinal effector promoting persistence within the microbiome and a major antigenic target of adaptive immunity, highlighting the balance between fitness and immune recognition required for persistent colonization.

Awp11 enhances fitness specifically in the context of a complete microbiota, though its mechanism remains unclear. Cluster V adhesins have been shown to mediate adherence to abiotic surfaces and cell aggregation (14), but their in vivo function is poorly defined. The best-characterized adhesins in *C. glabrata*, the EPA family, promote interactions with ovarian and intestinal epithelial cells, and at least one EPA adhesin plays a role in intestinal colonization (35–37). However, our data indicate that Awp11 is not necessary for adhesion to the intestinal epithelium. Rather, its role in enhancing colonization of the gut is dependent on the microbiome. In competition experiments, Awp11 enhanced colonization only in the presence of a complex microbiota, pointing to a role in inter-microbial interactions rather than host cell binding. Notably, a *C. albicans* Awp11 ortholog, Hyr1 promotes physical interactions with bacteria (38) and can also influence neutrophil activation (39), suggesting multiple potential mechanisms. Together, these data support a model in which Awp11 promotes competitive fitness within a polymicrobial environment rather than acting as a classical adhesin.

Our findings also reveal a functional interplay between Awp11 and mucosal immunity. Although *C. albicans* cell wall antibody targets have been well described (7, 9, 25, 40), very little is known about serological protein antigens of *C. glabrata*. Awp11 represents a protein antigen capable of eliciting robust B and T cell responses. IgA targeting of *C. albicans* cell wall effectors modulate pathogenesis and commensalism (7–9). Likewise, we show that IgA targeting of Awp11 reduces fungal fitness in the presence of a complex microbiota, suggesting a synergistic relationship between adaptive immunity and microbial competition. This parallels recent findings from Lentsch et al. and Santiago-Borges et al., where IgA-mediated protection against bacterial pathogens required competing bacterial species that occupy similar metabolic niches within the gut (41, 42). Importantly, our work extends this concept to cross-kingdom interactions, demonstrating that bacterial competition and antifungal IgA together shape fungal niche occupancy. In addition, Awp11 expression is associated with Th17 responses, although it remains unclear whether Awp11 itself is the dominant T cell antigen or indirectly promotes immune activation through its role in colonization.

A key feature of Awp11 is its dynamic regulation in most strains. Like other subtelomeric adhesins, *AWP11* expression in CBS138 is controlled by the Sir complex (32–34), though our data suggest additional layers of regulation. While a *sir3*Δ mutation increases *AWP11* transcription in CBS138 to those found in Cg27, it only partially enhanced recognition by Cg27-induced IgA.

Moreover, although overexpression of *AWP11* in CBS138 was sufficient to drive surface localization in vitro, this did not translate to increased IgA induction in vivo, indicating that environmental cues in the gut may constrain expression or protein display. Accordingly, increased transcription under host-like conditions did not consistently correlate with IgA binding, suggesting post-transcriptional regulation or altered trafficking.

Strikingly, *AWP11* expression varied depending on host immune status, with increased expression observed in immunodeficient settings. This supports the idea that epigenetic regulation of subtelomeric adhesins in *C. glabrata* may facilitate evasion of antigen-specific immunity (43), analogous to antigenic variation mechanisms in protozoan parasites (44). Our findings also raise the possibility that IgA itself, or other immune-associated factors, acts as a signal to modulate expression. Alternatively, heterogeneity in Awp11 expression within a strain may allow IgA to select for subpopulations with expressing lower levels of this effector. Regardless of mechanism, these findings suggest a form of antigenic modulation in response to host immunity.

These observations open several avenues for future investigation. Awp11 represents a promising candidate for therapeutic targeting, particularly as a vaccine antigen aimed at limiting intestinal reservoirs of *C. glabrata* that can seed systemic infection. The requirement for a complex microbiota in IgA-mediated fitness costs suggests that combining Awp11-targeted strategies with microbiota-based interventions, such as probiotics, may enhance efficacy. It will also be important to investigate how IgA influences susceptibility to antifungal drugs in the intestine, a site that promotes the acquisition of antifungal resistance in *C. glabrata* (45). However, it remains unclear whether Awp11 plays a similar role in human gut colonization or elicits comparable immune responses. Addressing these translational gaps will be critical for evaluating the clinical relevance of Awp11-targeted approaches.

## Materials and Methods

### Fungal Strains, Media, and Reagents

Fungal strains were grown overnight at 30 °C in yeast peptone dextrose (YPD) medium unless otherwise indicated. Inoculums were prepared by washing overnight cultures twice with PBS and resuspending cells to the indicated concentration for each assay. For formalin fixation, fungi were incubated in 10 mL of formalin for 10 minutes, followed by three washes with 10 mL PBS.

For antibiotic-treated experiments, sterile drinking water was supplemented with 50 µg-mL^-1^ each of ampicillin (RPI), neomycin (RPI), and gentamicin (CHEM-IMPEX). NAT-resistant transformants were selected by supplementing YPD with nourseothricin (RPI) at a final concentration of 200 mg-mL^-1^.

URA3 auxotrophs were selected on synthetic complete (SC) medium composed of 0.17% yeast nitrogenous base without amino acids or ammonium sulfate (RPI), 0.2% SC amino acid mixture (MP Biomedicals), and 2% glucose, supplemented with 100 µg-mL^-1^ 5-fluoroorotic acid (5-FOA). URA3 prototrophs were selected on SC–URA dropout medium containing 0.17% yeast nitrogenous base, 0.2% SC–URA mixture (MP Biomedicals), and 2% glucose.

To activate CRISPR mutagenesis, cells were grown in SC–CYS–MET medium consisting of 0.17% yeast nitrogenous base, 2% glucose, and individually added amino acids at concentrations equivalent to those in the SC amino acid mixture, excluding cysteine and methionine (18).

### Strain creation

Fungal strains used in this study are listed in Table S5. *URA3*𝛥 strains in the CBS138 and Cg27 background were generated as described (46). We utilized a CRISPR-Cas9 system as described in (18) by inserting an *AWP11* gRNA into pRS470 (pLH012). A Cg27 *URA3*𝛥 was then transformed with pLH012 and selected for on URA dropout media. The CRISPR Cas9 system was activated by overnight growth in methionine and cysteine deficient synthetic complete media. Mutations in *AWP11* were verified whole genome Illumina sequenced with SeqCenter. The plasmid pLH012 was cured from candidate colonies by overnight growth and counter selection on 5-FOA media. *URA3*𝛥 *ChrH::URA3* complement strains were generated in the CBS138 and Cg27 background as well as *awp11JDmut^fs^ URA3*𝛥 *ChrH::URA3* in the Cg27 background, by transforming via a homologous recombination fragment taken from pKO22. *URA3* integration was confirmed using primers 507 and 512.

To create a second loss of function mutant in the Cg27 background and the first in the CBS138 background each corresponding *URA3*𝛥 strain was transformed via a homologous recombination fragment taken from pLH016, then selected for on URA dropout media. *URA3* integration into *AWP11* was confirmed using primers 631 and 633. The result being Cg27 *awp11JD*𝛥*::URA3* and CBS138 *awp11*𝛥*::URA3*.

To constitutively express *AWP11* in CBS138 the WT CBS138 background was transformed via a homologous recombination fragment taken from pLH020, then selected for on NAT media. Integration of NAT and the Tef1 promoter into the native *AWP11* locus was confirmed using primers 334 and 371. The result being CBS138 pTEF1-*AWP11*.

To constitutively express OVA-GFP in the Cg27 and Cg27 *awp11JDmut^fs^* backgrounds the URA3 auxotroph of each background were transformed via a homologous recombination fragment taken from pBH002 using XhoI and HindIII restriction enzymes, then selected for on URA dropout media. Verification of OVA-GFP expression was determined with primers 517 and 518. Integration into chromosome M was confirmed using primers 484 and 524. The result being Cg27 *URA3*𝛥 *ChrM::URA3-OVA-GFP* and Cg27 *awp11JDmut^fs^ URA3*𝛥 *ChrM::URA3-OVA-GFP*.

Primer sequences located in Table S4.

### Vector creation

Vectors used in this study are listed in Table S6. pLH020 and pLH016 were generated by the Genscript FLASH gene service. The pKO22 plasmid, used to restore URA3 to auxotrophs, was generated using the NEB builder HiFi DNA assembly cloning kit with 3 fragments and a pUC19 backbone cut with XbaI restriction enzyme. The left and right chromosome M neutral site homology fragments were amplified using primers 507/508, and 511/512 respectively with Cg27 genomic DNA. The URA3 fragment was amplified using primers 509 and 510 on plasmid pCU-MET3-Hermes.

The pBH002 plasmid, used for expressing OVA-GFP, was generated by HiFi assembly of one fragment in the pKO22 background cut with NotI restriction enzyme. The OVA-GFP fragment was generated using primers 517 and 518 on plasmid pOJ002.

The pLH012 plasmid, used to induce CRISPR Cas9 mediated indels, was generated by HiFi assembly replacing the sgRNA sequence on pRS470 with an *AWP11* specific sgRNA fragment. This fragment was acquired in the form of a duplex primer ordered from IDT and is listed in Table S4 as 482.

### Mice

WT C57BL/6J, μMT^-/-^ (B6.129S2-*Ighm^tm1cgn^*/J), and *Tcrb^-/-^d^-/-^* (B6.129P2-*Tcrb^tm1Mom^Tcrd^tm1Mom^*/J) mice were obtained from The Jackson Laboratory (Bar Harbor, ME). B6.IgA^-/-^ (IgA^-/-^) were a gift from Dr. June Round PhD (University of Utah). For studies comparing WT and μMT^-/-^ or IgA^-/-^mice, these mice were bred in the University of Colorado Anschutz AMC Animal Facility under specific-pathogen free conditions and co-housed for 2-3 weeks post weaning. For WT B6 only and *Tcrb^-/-^d^-/-^* (TCRKO) experiments, mice were obtained directly from The Jackson Laboratory before the start of the experiment. C57BL/6J GF mice were bred in sterile isolator bubbles by the University of Colorado Anschutz Gnotobiotic core. Experimental mice were transferred to individually ventilated cages (IVC) (Allentown Inc) and treated with broad spectrum antibiotics (50 μg-ml^-1^ ampicillin, neomycin and gentamycin) in the drinking water to maintain sterility. The competition between Cg27 and Cg27 *awp11JD^fs^* in GF mice (Fig. 4C) was performed without added antibiotics. Prior to experimental use, GF mouse stool was tested for bacterial and fungal contamination using 16S and ITS2 PCR. Both male and female mice were used for these studies. All mice were between 6-12 weeks of age, were fed a standard diet. All studies were approved and performed in compliance of the by the University of Colorado Anschutz Institutional Animal Care and Use Committee (protocol 01221).

### Mouse Intestinal Colonization Models

For the clinical isolate screen, 34 strains were assigned to one positive control (Cg1) and five pools (Pool 1: Cg2–7; Pool 2: Cg8–13; Pool 3: Cg14–20; Pool 4: Cg21–27; Pool 5: Cg28–34). Pools were combined in equal ratios at a final concentration of 10^8^ CFU-ml^-1^. Mice were treated with antibiotics in their drinking water and then gavaged 3x weekly with 100 µL of corresponding group. Cg5 was removed due to a growth defect in YPD. Fecal samples were collected at days 7, 14, and 21 post-initial gavage. Mice were euthanized four weeks after colonization, and small intestinal, cecal, and colonic contents were collected.

For mono-colonization experiments, mice were intestinally colonized by oral gavage using prepared inoculums. Gnotobiotic mice and all antibiotic-treated mice received antibiotics in drinking water for 3–7 days prior to gavage, whereas gnotobiotic mice in Figure 4 did not receive antibiotics. Unless otherwise indicated, mice were colonized for 21 days and then euthanized for collection of intestinal contents or tissues. Gnotobiotic and antibiotic-treated mice received a single gavage of 100 µL inoculum at 10^7^ CFU-ml^-1^. SPF mice received either a single gavage or gavage three times weekly with 100 µL inoculum at 10^8^ CFU-ml^-1^, or an equivalent formalin-fixed inoculum.

### In vivo fungal competition

*C. glabrata* Cg27 versus Cg27 *awp11JD^fs^* or *awp11JD*Δ and CBS138 versus CBS138 *awp11*Δ expressing NEON, iRFP670 or no fluorescent reporter were mixed 1:1 and 10^6^ (antibiotic-treated) or 10^7^ (SPF) total cells were orally gavaged into mice. To control for the fitness impacts associated with the fluorescence marker, each competition (except Fig. 6E & 6F) was performed in duplicate with reciprocal expression of the fluorescent maker in each strain. For the fecal microbiota transplant (FMT) experiments (Fig. 5), Cg27 and versus Cg27 *awp11JD^fs^* expressing NEON were mixed 1:1 and 10^6^ CFU were gavaged into antibiotic treated mice. At day 28 day, antibiotics were removed and mice were placed on sterile water for 4 days. Mice were then given an FMT by gavaging with 100 mg-ml^-1^ homogenized stool pellets in anaerobic-reduced PBS from SPF B6 mice. For all experiments, stool samples were processed as follows. Stool samples (200-500 mg), collected at indicated timepoints, or endpoint intestinal contents and tissue were homogenized in PBS, and plated onto YPD+ penicillin and streptomycin plates. Plates were scraped and resuspended in 5 ml PBS and an aliquot was used to quantify NEON or iRFP670 vs non-fluorescent frequency by flow cytometry on the Becton Coulter CytoFlex. Competitive index (CI) was calculated using the following formula: % NEON or iRFP+ recovered / % inoculum NEON or iRFP+.

### Intestinal colonization-intravenous dissemination model

Intestinal colonization of antibiotic treated mice was performed as described above. B6 WT and μMT^-/-^ mice were gavaged on day 1 with 10^6^ CFU Cg27 or CBS138. In another set of experiments, antibiotic treated B6 mice were gavaged with Cg27, CBS138, or PBS control for 21 days. On day 21 post-colonization, mice were anesthetized with isoflurane and retro-orbitally injected with 10^7^ CFU CBS138 or Cg27i n 0.1 ml sterile PBS. Mice were monitored daily for weight loss or signs of physical distress. 7 days post infection mice were euthanized, and spleens, kidney, and liver were, weighed, and homogenized (bead beat for 30 S in 1 ml PBS) and dilutions were plated on YPD + Penicillin/Streptomycin media to quantify CFUs. CFUs were normalized to input tissue weight.

### Transcriptional analysis of in vivo or in vitro fungi

Cecal contents (∼500 mg) were collected from mice and immediately frozen on dry ice and transferred to -80°C. For in vitro RNAseq, overnight cultures were diluted 1:100 in YPD and grown for 6 hours shaking at 30 °C. Cultures were then spun down at 3000 RPM x 5 minutes, and the pellet was isolated for RNA extraction. Fungi and cecal contents were lysed by bead beating 3 x 30 sec with 0.5 mm glass beads (BioSpec) in RPE buffer + Beta-Mercaptoethanol. RNA was isolated using the RNeasy Mini kit with DNAse I treatment (Qiagen). For qRT-PCR, cDNA was synthesized using qScript cDNA synthesis kit (Quanta Biosciences), and transcripts were measured using PowerUp SYBR Green Master Mix (Applied Biosystems) and following primers: *AWP2* (348/349), *AWP4* (380/381), *AWP8* (372/373), *AWP10* (368/369), *AWP11* (370/371) (Table S4). Samples were normalized using the housekeeping gene *ACT1*. Targeted qPCR was analyzed on the QuantStudio 6 Real-Time PCR System.

For RNA sequencing, the Novogene eukmRNA-seq library prep kit was used to generate libraries. Libraries were sequenced on an Illumina [platform, e.g. NovaSeq 6000] using paired-end 150 bp reads (2×150 bp). All paired-end reads were assessed for quality before and after trimming using FastQC v0.11.9 (47). The Illumina universal sequencing adapters were trimmed and low-quality reads removed (-q 30 --minimum-length=10 --pair-filter=any) using cutadapt v4.2 (48). All reads in each sample were screened for mycoplasma contamination using FastQ Screen v0.16.0 (49). Read pairs were aligned to the allele A haplotype of the *C. glabrata* genome and counts quantified (--quantMode TranscriptomeSAM) using STAR v2.7.10b with the following parameters specified: --outFilterScoreMinOverLread 0.40 --outFilterMatchNminOverLread 0.40 (50). Quality was assessed by collecting metrics using the Picard suite of tools v2.27.5 and MultiQC v1.14 to summarize metrics from upstream processing (51, 52). Genes were filtered prior to differential expression analysis, retaining only those with a minimum of 5 reads in at least 50% of samples within a biological group and present across at least 2 comparison groups. PCA was performed to identify technical and biological sources of variation in the dataset. PC3 showed the strongest correlation with batch and was therefore included as a covariate in the GLM. Normalization and differential expression analysis were performed using DESeq2 (53). To account for large log2(FC) values for genes where dispersion estimates may be high due to low gene counts or larger variance within biological replicates, the apeglm shrinkage estimator was applied to get a more realistic conservative estimate of true log2(fc) values (53). The top 50 most differentially regulated genes were selected by sorting by adjusted p-values < 0.05 and were ranked by absolute log2 fold change. Hierarchal clustering in the heatmaps was performed using correlation distance comparing gene expression between samples.

### *C. glabrata* cell wall isolation and western blot

Overnight cultures were pelleted, resuspended in PBS, and then centrifuged at maximum speed at 4 °C. Supernatants were removed, pellets were flash-frozen on dry ice, and resuspended in 600 µL PBS supplemented with 0.5 ml 0.5 mm glass beads and 1X Protease inhibitor (Roche). Cells were lysed by bead beating (30 s, 1 min on ice) for 10 cycles. Lysates were transferred to pre-weighed tubes after bead settling, and residual protein was recovered by washing beads twice with 400 µL 1 M NaCl. Clarified lysates were centrifuged at maximum speed for 10 min at 4 °C, resuspended in 1 mL 1 M NaCl, and inverted for 5 min; this wash was repeated five times, followed by a final wash in autoclaved water. Protein concentrations were determined by BCA and normalized to 1 mg-mL^-1^. Samples were mixed with 4× Bolt LDS sample buffer and 10× Bolt reducing agent (Thermo) to a final volume of 100 µL, boiled at 95 °C for 5 min, and separated by SDS-PAGE using the Mini Blot Module and protocol (Invitrogen). Following transfer, blots were blocked in 5% milk for 1 hr and then incubated overnight with SI wash from uncolonized or mono-colonized GF mic at 1:40 dilution in PBS. Membranes were washed in PBS-0.05% Tween20 (PBS-T) fand then incubated with anti-mouse IgA-HRP (Southern biotech) diluted 1:2000 in PBST for 2 hrs. Membranes were analyzed by chemiluminescence using Super Signal West PICO Plus (Thermo).

### In-gel digestion for proteomic analysis

Electrophoretic protein bands were excised from Coomassie-stained gels, destained, and subjected to in-gel reduction, alkylation, and overnight trypsin digestion as previously described (54). Following overnight digestion, samples were acidified with 5% formic acid (FA), and tryptic peptides were extracted in 30 µl of 50% acetonitrile, 1% FA. Digests were dried in a vacuum centrifuge and resuspended in 0.1% FA.

### LC-MS/MS analysis

Liquid chromatography-tandem mass spectrometry (LC-MS/MS) was performed using an Easy nLC 1000 instrument coupled with a Q Exactive™ HF Mass Spectrometer (both from ThermoFisher Scientific). For each sample, 3 μg of digested peptides were loaded on a C18 column (100 μM inner diameter x 20 cm) packed in-house with 2.7 μm Cortecs C18 resin, and separated at a flow rate of 0.4 μl-min^-1^ with solution A (0.1% FA) and solution B (0.1% FA in ACN) under the following conditions: isocratic at 4% B for 3 min, followed by 4%-32% B for 102 min, 32%-55% B for 5 min, 55%-95% B for 1 min and isocratic at 95% B for 9 min. Mass spectrometry was performed in data-dependent acquisition (DDA) mode. Full MS scans were obtained from m/z 300 to 1800 at a resolution of 60,000, an automatic gain control (AGC) target of 1x10^6^, and a maximum injection time (IT) of 50 ms. The top 15 most abundant precursors with an intensity threshold of 9.1 x 10^3^ were selected for MS/MS acquisition at a 15,000 resolution, 1 x 10^5^ AGC, and a maximum IT of 110 ms. The isolation window was set to 2.0 m/z and ions were fragmented at a normalized collision energy of 30. Dynamic exclusion was set to 20 s.

### Global proteomic data analysis

Data was searched using MSFragger v4.1 via FragPipe v22.0 (55). Precursor tolerance was set to ±15 ppm and fragment tolerance was set to ±25 ppm. Data was searched against UniProt restricted to the *Candida glabrata* reference proteome (UP000002428) with added common contaminant sequences (56) as well as two predicted amino acid sequences representing copies of the CAGL0J12067 gene identified via genome sequencing of this *C. glabrata* strain (5,309 total sequences). Enzyme cleavage was set to semi-specific trypsin for all samples allowing for 2 missed cleavages. Fixed modifications were set as carbamidomethyl (C). Variable modifications were set as oxidation (M) and acetyl (protein N-terminus). Label free quantification was performed using IonQuant v1.10.27 with match-between-runs enabled and default parameters. Results were filtered to 1% FDR at the peptide and protein level. For variant analysis of the CAGL0J12067 gene product, peptides unique to each copy based on the predicted amino acid sequences used in database searching were assessed to determine copy-specific expression.

### Antibody reactivity to in vitro grown fungi and Total IgA ELISA

Intestinal contents were weighed and resuspended at 100 mg-ml^-1^ in sterile PBS. Following homogenization, samples were spun at 3,000 RPM for 10 minutes and supernatant was isolated for antibody reactivity. Serum was extracted from whole blood following submandibular bleeds or cardiac puncture at experiment endpoint. Intestinal contents diluted 1:1 and serum diluted 1:100 in PBS. Antibody reactivity to in vitro grown fungal strains was performed as described in (9), and analyzed by flow cytometry using the LSR Celesta or Beckman Coulter CytoFlex and FlowJO. Total IgA ELISAs (Invitrogen) were performed according to manufactures protocol.

### Quantification of IgA binding and cell wall components in vivo

IgA quantification and cell wall component analysis was adapted from (9). In brief, intestinal contents were isolated from SI, cecum, or colon (stool) and resuspended at 100 mg-ml^-1^ sterile PBS by manual disruption. Samples were centrifuged at 100 G for 30 sec to pellet debris, and then 50 μl of supernatant was added to 96 well U-bottom plate. Samples were spun at 3,000 RPM x 5 min, washed 2X with FACS buffer (PBS + 2% FBS + 1 mM EDTA), and then blocked with 10% FBS in PBS (IgA, Concavalin A, CalcoFluor White), or 1% BSA-5mM EDTA-3% goat Serum (Fc-Dectin1) for 20 minutes at RT. For IgA quantification samples were stained in 50 μl FACS buffer containing 1:250 anti-IgA-PE (eBioscience clone mA-6E1) and 1:250 anti-Candida-FITC (Meridian Biosciences) for 20 minutes at RT. For ConA and CFW quantification, samples were stained with 50 μl FACS buffer with 25 μg-ml^-1^ ConA-APC, 2 μg-ml^-1^ CFW, and 1:250 anti-Candida-FITC for 30 min at 4°C. Fc-hDectin1 (Invivogen) was diluted in 3% goat serum in PBS at 5 μg-ml^-1^ for 60 min at 4°C, washed 2X and then stained with 50 μl 1:500 anti-human IgG AF594 (Jackson labs) and 1:250 anti-Candida-FitC in FACS buffer for 20 min at 4°C. All samples were then washed in FACS buffer, fixed in 2% paraformaldehyde (Sigma) for 10 minutes and analyzed on Beckman Coulter CytoFLEX.

### Lamina propria, MLN, and PP preparation and flow cytometry analysis

The small intestine was excised, and Peyer’s patches and fat were removed from the entire SI. The distal 1/3 of the SI (ileum) was then fileted open longitudinally and cut into four equal pieces for lamina propria isolation using a protocol adapted from (57). Briefly, SI pieces were incubated two times in PBS containing 2% FBS, 5 mM EDTA (Fisher), and 1 mM DL-Dithiothreitol (DTT) (Thermo) for 15 min shaking at 37°C. Samples were washed briefly in PBS and then digested in 10 mls prewarmed R10 media (RPMI (Gibco), 10% FBS (VWR, 1 mM HEPES (Fisher), 1X Penn/Strep (Cytiva)) containing 0.5 mg/ml Collagenase D (Roche), and 0.5 mg/ml DNaseI (Sigma) for 40 min at 37°C shaking. Cell suspensions were briefly vortexed and passed through a 40 μM filter. Tissue remaining in filter was gently grinded using a syringe filter and 20 mls of ice cold FACS buffer was added. Live immune cells were isolated using a 40:70% Percoll gradient (Fisher). Single cell suspensions of PP and MLN were isolated by grinding through 40 μM filters into R10 media.

For immune cell stimulation, 1-2 x 10^6^ cells were added to a 96 well U bottom plate and incubated in R10 media with 200 ng/ml Phorbol 12-myristate 13-acetate (PMA), 1 μg-ml^-1^ Ionomycin (Sigma), and 1X GolgiPlug (BD) for 4 hours 37°C 5% CO_2_. Following stimulation, wells were washed 1X with FACS buffer and then labeled with Live/Dead Fixable Blue (Thermo) for 15 minutes. Samples were subsequently incubated with TruStain Fc block (Biolegend) for 10 minutes and then the following anti-mouse antibodies in Brilliant Stain buffer (BD): anti-CD3-BUV395 (Clone 145-2C11), anti-CD4-BUV395 (clone GK1.1), anti-CD45-BUV805 (clone 30-F11), anti-IgM-RB780 (clone II/41), anti-CD8a-BUV615 (clone 53-6.7) (BD Biosciences), anti-B220-SparkViolet538 (clone RA3-6B2), anti-CD95-APC (clone SA367168), anti-CD138-PE-Fire810 (clone 281-2), anti-CXCR5-BV650 (clone L13D7), anti-PD1-BV785 (clone 29F.1A12), anti-TACI-PE (clone 8F10), anti-GL7-Pacific Blue (clone GL7), and anti-IgD-BV605 (clone 11-26c.2a) (Biolegend). Samples were washed in FACS buffer and fixed overnight at 4°C using the FoxP3 Transcription Factor Staining Buffer kit (eBioscience). Cells were washed 2X in Permeabilization buffer and then stained with the following intracellular antibodies: anti-Foxp3-R718 (clone FJK-16S), anti-Tbet-PE-Cy5 (clone 4B10), anti-RORγT-PE-CF594 (clone Q31-378) (BD Biosciences), anti-IgA-FITC (clone mA-6E1, Invitrogen), anti-IL-17A-PE-Cy7 (clone TC11-18H10.1), and anti-IFNγ-BV421 (clone XMG1.2) (Biolegend). 123Count eBeads (Invitrogen) were added for cell quantification. These data were collected on the Cytek Aurora and analyzed using FlowJo.

### Caco-2 epithelial adhesion assay

Low-passage (<40) Caco-2 (BBE1) cells were counted by trypan blue exclusion and resuspended in DMEM supplemented with 10% FBS at 1 × 10^5 cells-ml^−1^. Cells were seeded into flat, tissue culture–treated 96-well plates and maintained at 37 °C with 5% CO₂. Media were replaced three times weekly until cells formed polarized monolayers (∼21 days). Overnight *C. glabrata* cultures were resuspended in tissue culture medium at 10^6^ cells-ml^−1^. Monolayers were washed twice with fresh medium to remove dead cells, followed by addition of 200 µL fungal suspension per well. Plates were centrifuged at 500 g for 1 min to promote cell–cell contact and incubated for 1 h at 37 °C with 5% CO₂. Non-adherent fungi were removed by two washes with medium, and cells were fixed with 2% paraformaldehyde. For fluorescence microscopy, wells were stained for 20 min in column buffer (HBSS + 0.5% FBS + 0.5 mM EDTA, + 1.0 mM HEPES) with DAPI (1:500) and anti-*Candida*-FITC (1:500). Wells were resuspended in 100 µL PBS for imaging. Images were acquired in triplicate per well, and adherent fungal and epithelial cells were quantified using automated fluorescence analysis.

### Analysis of Awp11 expression in cecal content growth assay

Cecal content culture conditions were adapted from Liang et al (31). Briefly, cecal contents from uncolonized IgA^-/-^ or germ-free mice were resuspended in sterile PBS at 25 mg-ml^-1^ for and filtered to remove debris. 200 μl of diluted contents per sample was added to a U bottom 96 well plate. Overnight cultures of *C. glabrata*, were washed 2X in PBS, enumerated, and normalized to 10^8^ cells-ml^-1^. For clinical isolate screen, all strains were normalized by OD600. 10 μl (10^6^ cells) was added to 96 well plate and incubated at 37°C 5% CO_2_ for 3 or 6 h. YPD control wells were included for all samples. Following incubation, RNA was extracted and *AWP11* qRT-PCR was performed as described above. Additional wells were used for analysis of Awp11 cell surface expression. Sample wells were washed 2X in FACS buffer and then incubated in 50 μl cecal content supernatant from germ free mice monocolonized with Cg27 (Awp11-specific IgA) for 1 h at 4°C. Wells were washed 2X in FACS buffer and then stained with 1:500 anti-Candida-Biotin (Meridian) for 20 min at 4°C and subsequently with 1:1000 Streptavidin-APC (Biolegend) and anti-IgA-PE (eBioscience) 20 min at 4°C. Samples were fixed in 2% PFA and analyzed by flow cytometry. IgA positivity was recorded as frequency IgA positive relative to Cg27 cecal content negative controls.

### Short-read whole genome sequencing

Genomic DNA was extracted from overnight YPD cell cultures gof 32 bloodstream isolates plus CBS138 using the MasterPure Yeast DNA extraction kit (Fisher). Sequencing libraries were prepared using the plexWell 384 kit (cat PW384, seqWell Inc, Beverly MA). Libraries were sequenced at Novogene Corporation Inc (Sacramento, CA) on a NovaSeq X Plus sequencer using a partial lane of a 25B flow cell in paired-end 150 bp mode. Reads were trimmed using fastp v1.1.0 (58) with default parameters. Trimmed reads were aligned to the CBS138 reference genome version s05-m03-r11 (https://www.candidagenome.org) using bwa-mem2 v2.2.1 (59) and secondary alignments were filtered out using samtools view v1.23 (60). Read duplicates were marked and removed using picard MarkDuplicates v3.1.1 (http://broadinstitute.github.io/picard). Read group tags were added using picard AddOrReplaceReadGroups v3.1.1. Variants were called using bcftools mpileup v1.19 (61) and bcftools call v1.19 with options -m -v. Single nucleotide variants of PHRED33 quality scores below 20 were excluded from the downstream analyses using bcftools view v1.19. Gene copy numbers were estimated using the per-position depth of coverage values obtained with samtools depth v1.23, normalized by the median depth of coverage of the whole genome in a custom Python v3.12 (62) script.

### Phylogenetic analysis

The phylogenetic tree comprising 32 bloodstream isolates and the type strain CBS138 was built from concatenated coding DNA sequence of 100 non-repetitive nuclear genes selected as follows. BLASTN v2.16.0 (63) was used to align each of the 5280 coding ORF DNA sequences (https://www.candidagenome.org) against all ORF sequences with parameters -word_size 11 - dust no -soft_masking false -perc_identity 95. The resulting alignments were analyzed using a custom Python v3.12 script. For each gene, a bit score threshold was defined as the minimum bit score for self-hits multiplied by 0.03. A given gene was selected as a candidate if at most one non-self hit had a bit score higher than the threshold. Out of 4216 genes meeting this criterion, 100 were selected at random and output in BED format. Consensus sequences were generated for each strain by applying single nucleotide variants to the reference genome sequence using bcftools v1.19. Consensus sequences were concatenated using a custom Python v3.12 script, resulting in a multiple sequence alignment of 229637 positions. A maximum likelihood phylogenetic tree was computed on the alignment using raxml-ng v1.1.2 (64) with the GTR+G model and 200 bootstrap trees. Phylogenetic tree visualization was done using FigTree v1.4.5 (http://tree.bio.ed.ac.uk/software/figtree/).

### Genomic DNA isolation for long-read sequencing

To prepare reads for Oxford Nanopore Minion sequencing, high molecular weight DNA was isolated from cells by the following method: cells from overnight cultures were pelleted and resuspended in digestion buffer (1M sorbitol; 0.1M EDTA pH 7.5, 0.5 mg-ml^-1^) 20T zymolyase) and incubated at 37℃ for 60 minutes. Cells were gently pelleted (4000 rpm for 1 minute), resuspended in lysis buffer (50 mM Tris, 20 mM EDTA 1% SDS) and incubated at 65℃ for 30 minutes. 5M potassium acetate was added to a final concentration of 1.4M and samples were incubated at 4℃ for 1hr. Precipitated debris was pelleted at 14000 rpm for 15 minutes and the supernatant retained. Nucleic acids were precipitated with the addition of isopropanol, recovered by pelleting, and washed with 70% ethanol. RNA was degraded from the nucleic acid pellet by incubation in RNase buffer (50 mM Tris, 20 mM EDTA, 0.15 mg-ml^-1^RNase A) at 65 ℃ for 15 minutes, followed by the addition of ammonium acetate (2.5M) and a second precipitation of the remaining nucleic acid with isopropanol. Pellets were washed with 70% ethanol and dissolved in water overnight at room temperature.

### Long-read sequencing and assembly

Oxford Nanopore long reads were obtained in two distinct batches. Batch #1 included strains Cg1, Cg14, Cg27 and CBS138, while batch #2 included strains Cg1, Cg14 and Cg27. High-quality genomic DNA was extracted from cell cultures as described above. For batch #1, libraries were made using the SQK-LSK110 kit (Oxford Nanopore Technologies, Oxford UK). Libraries were sequenced individually on a GridION X5 Mk1 sequencer using four R9.4.1 flow cells (cat FLO-MIN106) at 450 bps for 48 hrs. Basecalling was run using Guppy v6.4.6 with the high accuracy model version 2021-05-17_dna_r9.4.1_minion_384_d37a2ab9. For batch #2, libraries were prepared in multiplex using the SQK-NBD114-24 kit. The library pool was sequenced on a MinION Mk1B sequencer with a single R10.4.1 flow cell (cat FLO-MIN114) at 400 bps for 72 hrs. Basecalling was run using Dorado v7.6.8 with the super accuracy model version dna_r10.4.1_e8.2_400bps_sup@v5.0.0.

Raw reads in FASTQ format were merged for batches #1 and #2 and used as input for de novo assembly with Canu v2.2 (65) with parameters genomeSize=13m -fast. Raw contigs were aligned against the CBS138 reference genome using MUMmer nucmer v4.0.0rc1 (66) with parameter --mum. Alignment coordinates were extracted using MUMmer show-coords v4.0.0rc1 with parameters -r -T- d. Contigs were reordered and reverse-complemented as needed using a custom Python v3.12 script. Chromosomal rearrangements were called from dot plots generated in a custom Python v3.12 script from alignment segments longer than 5 kb. The presence of the J-D translocation in Cg20 and Cg4 was called by aligning the corresponding Illumina short reads as described above, using our Cg27 de novo assembly as a reference sequence and visualizing the pileups JBrowse v4.1.3 (67).

### Statistical analysis

All statistical analysis were performed using Prism version 10 (GraphPad Software). Specific statistical tests are noted in the figure legends. All graphs were produced using Prism or R (Vienna, Austria) and the ggplot2 graphing package (68).

## Data Availability

RNA and DNA sequencing data is available under BioProject PRJNA1459241. Code used for analyzing *C. glabrata* genomes are available: https://github.com/mhenault1/glabrata_adhesins

## Acknowledgements

We thank the University of Colorado Anschutz Gnotobiotic Core, Proteomics Core, and Cancer Center Flow Cytometry Core for their expert technical support and resources that were essential to this work. We are grateful to Rebecca Shapiro PhD and Laetitia Maroc PhD for providing CRISPR plasmids, Kyle Cunningham PhD for valuable guidance on URA knockout protocols, and Kim Hanson MD for generously supplying *Candida glabrata* clinical strains.

## Author contributions

Conceptualization, L.H, O.J, K.S.O.; Data curation, L.H, O.J., M.H, L.R.H; Formal analysis L.H, O.J, M.H, M.M.; Investigation, L.H, O.J, E.T, C.B, B.E.H. K.S.O, M.H, L.R.H., B.C.R., M.M.; Project administration, K.S,O; Writing – review and editing, all authors.

## Supplemental information

Document S1. Figures S1-S10

Table S1. Fig. S1 Statistics

Table S2. Differential expression analysis

Table S3. *C. glabrata* cell wall proteomics.

Table S4. Primers

Table S5. Fungal Strains

Table S6. Plasmids

