## Supplementary Figures for "A *Candida glabrata* adhesin-like effector drives fitness and immunogenicity in the gut"

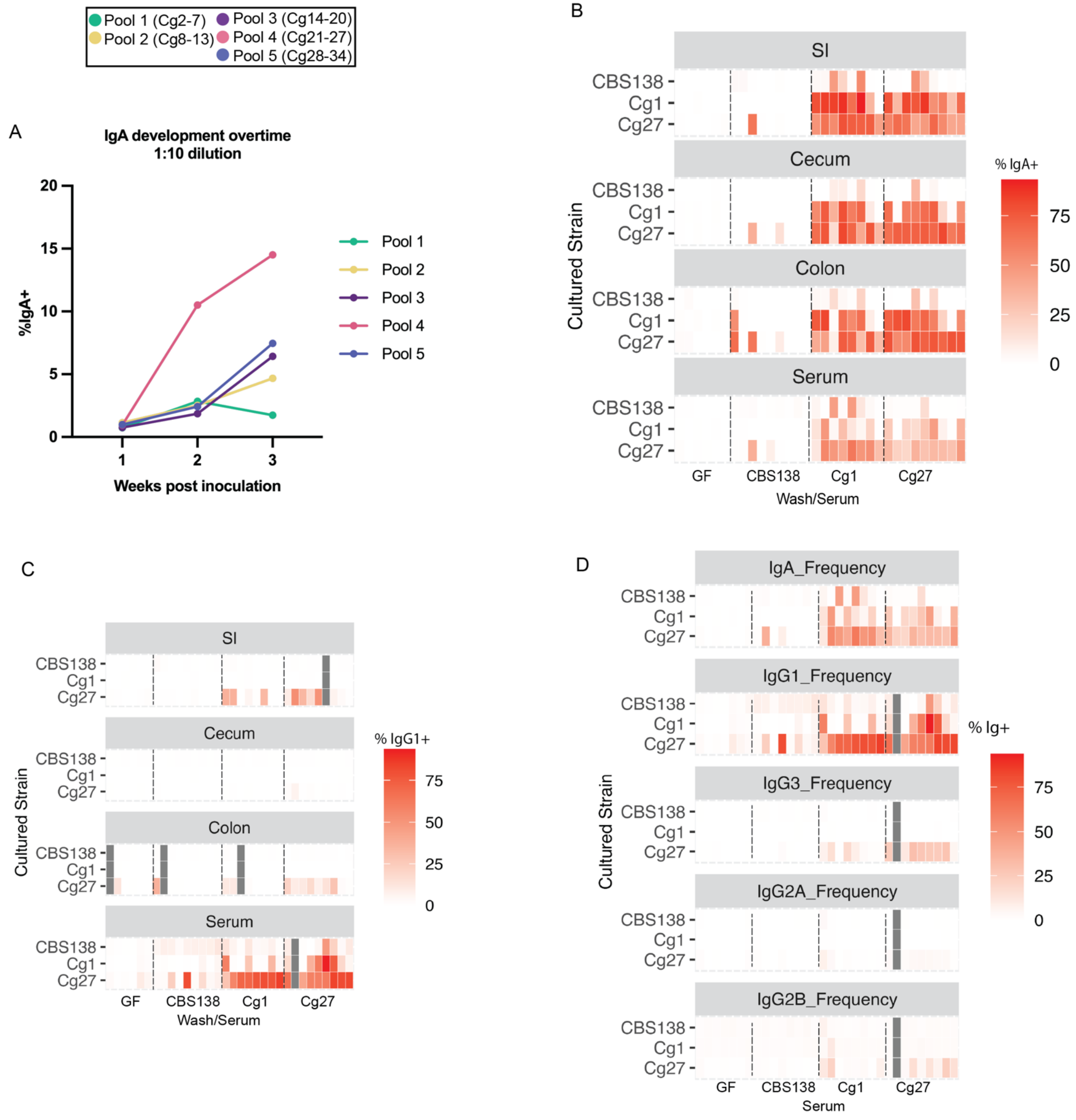


Fig. S1. ***C. glabrata* strain specific mucosal and systemic immunoglobulin induction in mono-colonized gnotobiotic mice.** (A) IgA binding from each pool's fecal wash incubated with its pool inoculum in vitro, measured as IgA bound frequency. Heatmap of day 28 SI, cecal, and colon wash and serum from gnotobiotic mice IgA (B) and IgG1 (C) binding frequency incubated with indicated cultured strains measured by flow cytometry. Each square represents an individual mouse. (D) Heatmap of day 28 serum immunoglobulin isotype binding to in vitro grown *C. glabrata*. Each square represents an individual mouse. (A-D) Two-way ANOVA with Tukey's multiple comparison test. n=5-9 mice per group Each dot represents an individual mouse across three independent experiments. Corresponding p values located in Table. S1.

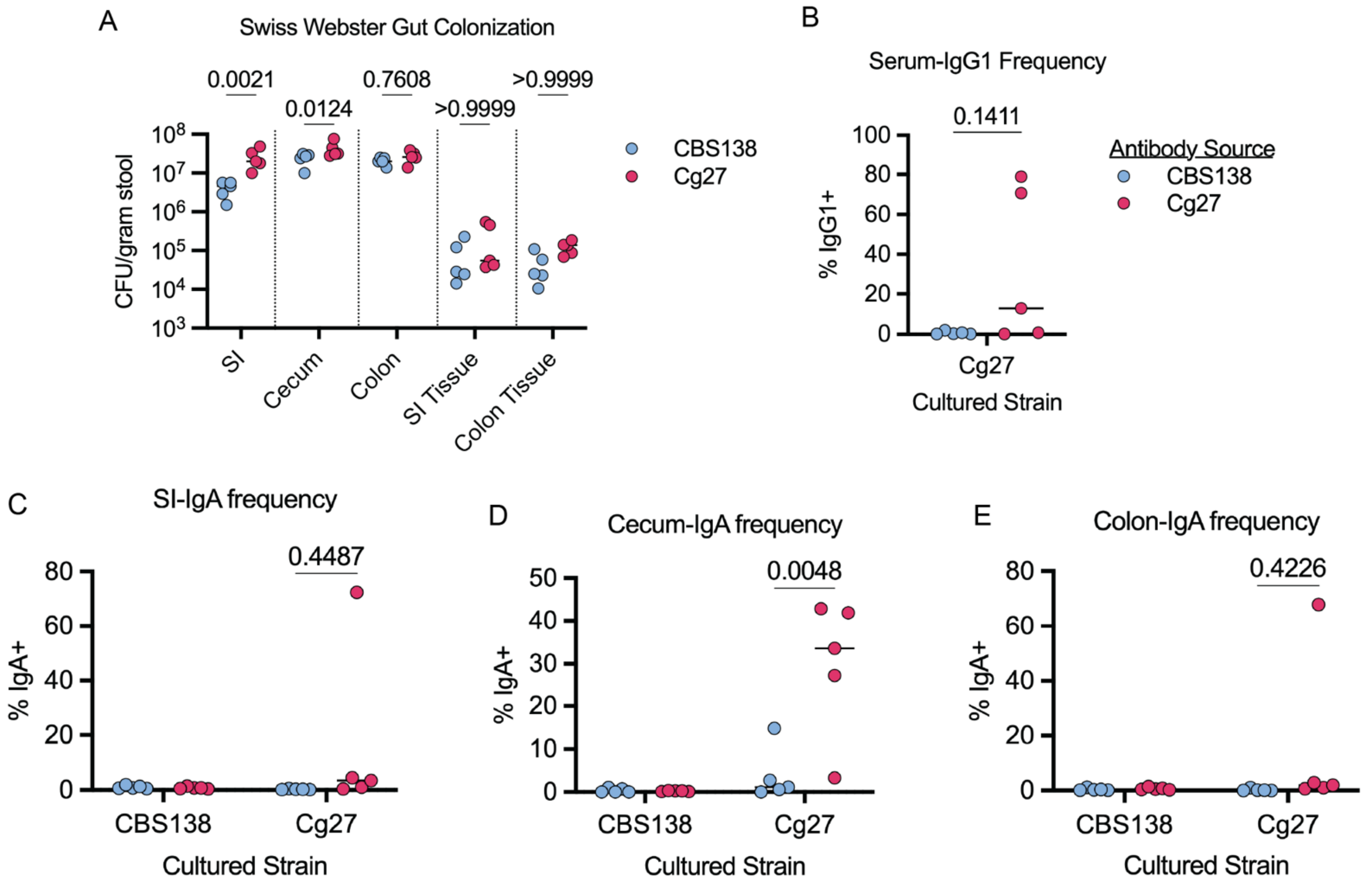


Fig. S2. ***C. glabrata* strain specific mucosal and systemic immunoglobulin induction in antibiotic treated Swiss Webster mice.** (A) Intestinal content and tissue fungal burden represented as CFU-gram$^{-1}$ contents/tissue. (B) Frequency of day 21 serum IgG1 binding to in vitro grown *C. glabrata* Cg27. Frequency of day 21 SI (C), cecal (D), and colon (E) content wash IgA binding to indicated *C. glabrata* strains. All p values calculated by Mann-Whitney U test. n=5 mice per group from one independent experiment. Each dot represents an individual mouse.

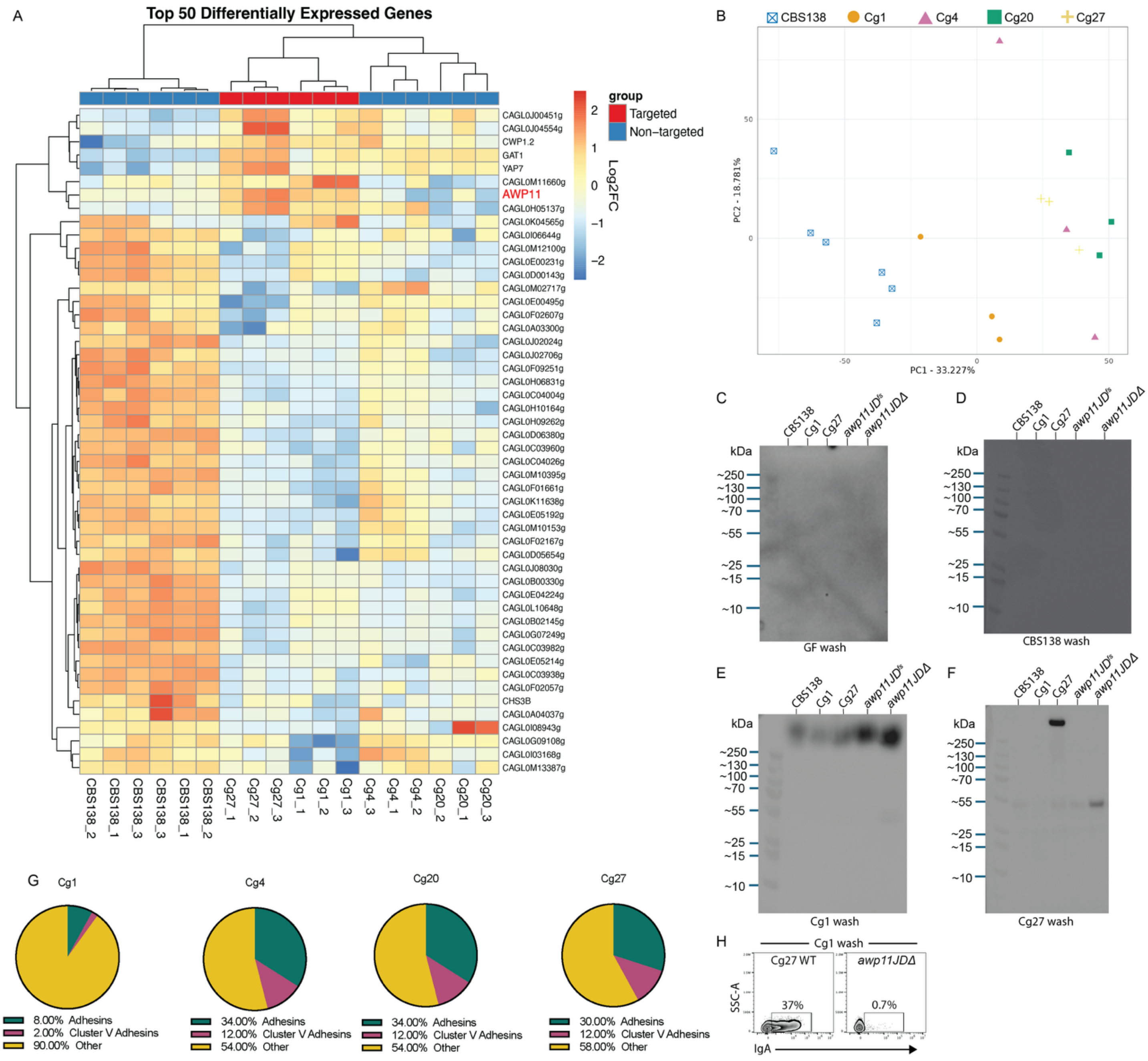

Fig. S3. **Transcriptional and cell wall western blot analysis of *C. glabrata* isolates.** (A) Top 50 differentially expressed genes comparing Cg1 and Cg27 (IgA Targeted) to CBS138, Cg4, and Cg20 (non-targeted). (B) *C. glabrata* in vitro RNA-seq PCA plot. Full length images of cell wall protein western blots blotted with GF (C), CBS138 (D), Cg1 (E), or Cg27 cleared intestinal wash (F). (G) Proportion of adhesins among the top 50 differentially expressed genes comparing Cg1, Cg4, Cg20, or Cg27 individually to CBS138. (H) Representative flow cytometry plots of IgA binding to Cg27 and *awp11Δ CBS138* grown in vitro following incubation with Cg1 monocolonized GF cleared intestinal wash.

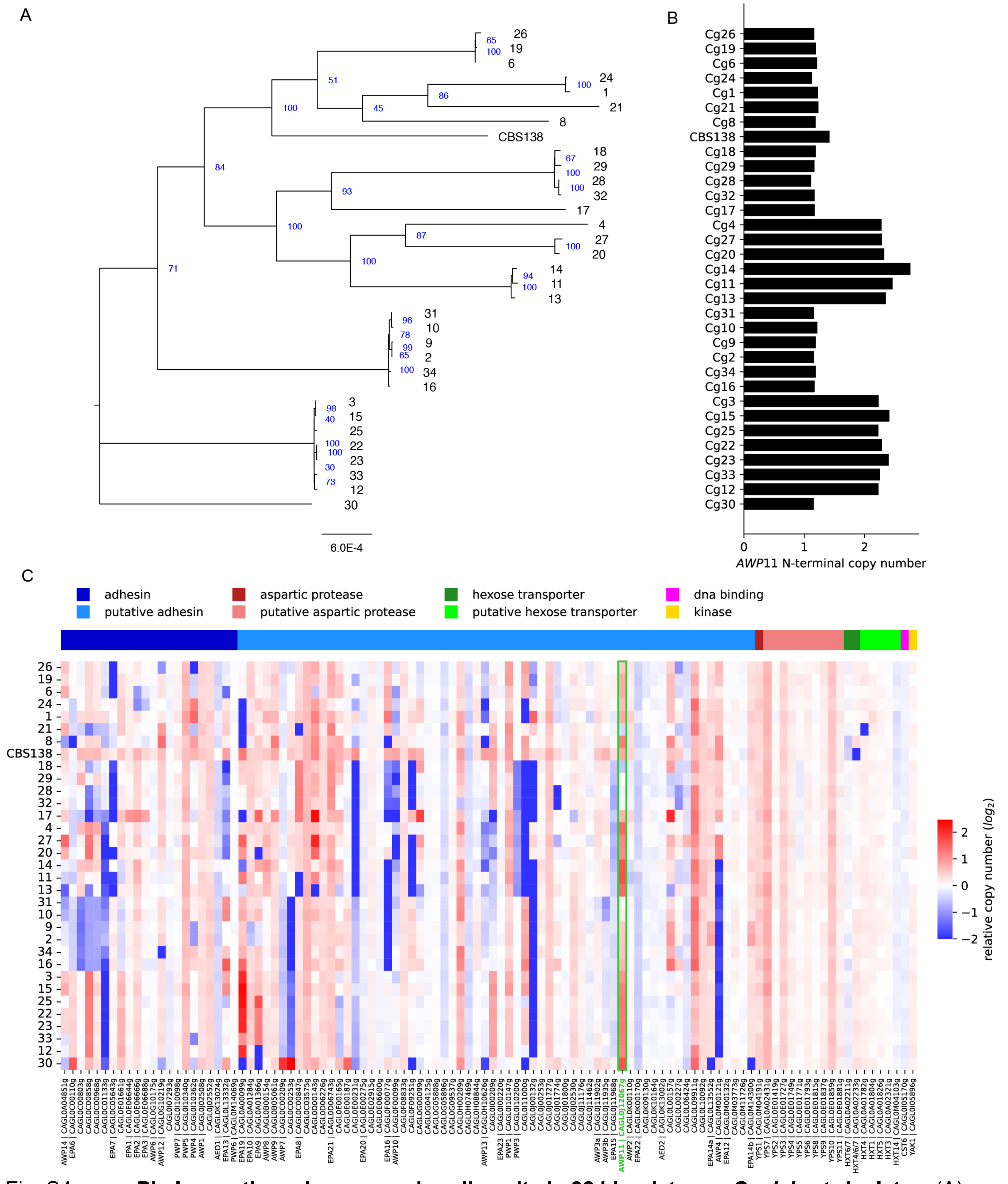


Fig. S4. **Phylogenetic and copy-number diversity in 32 bloodstream *C. glabrata* isolates.** (A) Maximum likelihood phylogenetic tree computed on 100 randomly selected single copy nuclear protein gene DNA sequences. Support values from 200 bootstrap trees are shown in blue. Scale is in substitutions per site. (B) Copy number variation of the 5' portion of *AWP11*. (C) Copy number variation for 106 genes comprising annotated adhesins and other pathogenicity-associated genes. *AWP11* is highlighted in green. Strain order on the Y axis is the same as for the phylogenetic tree in (A).

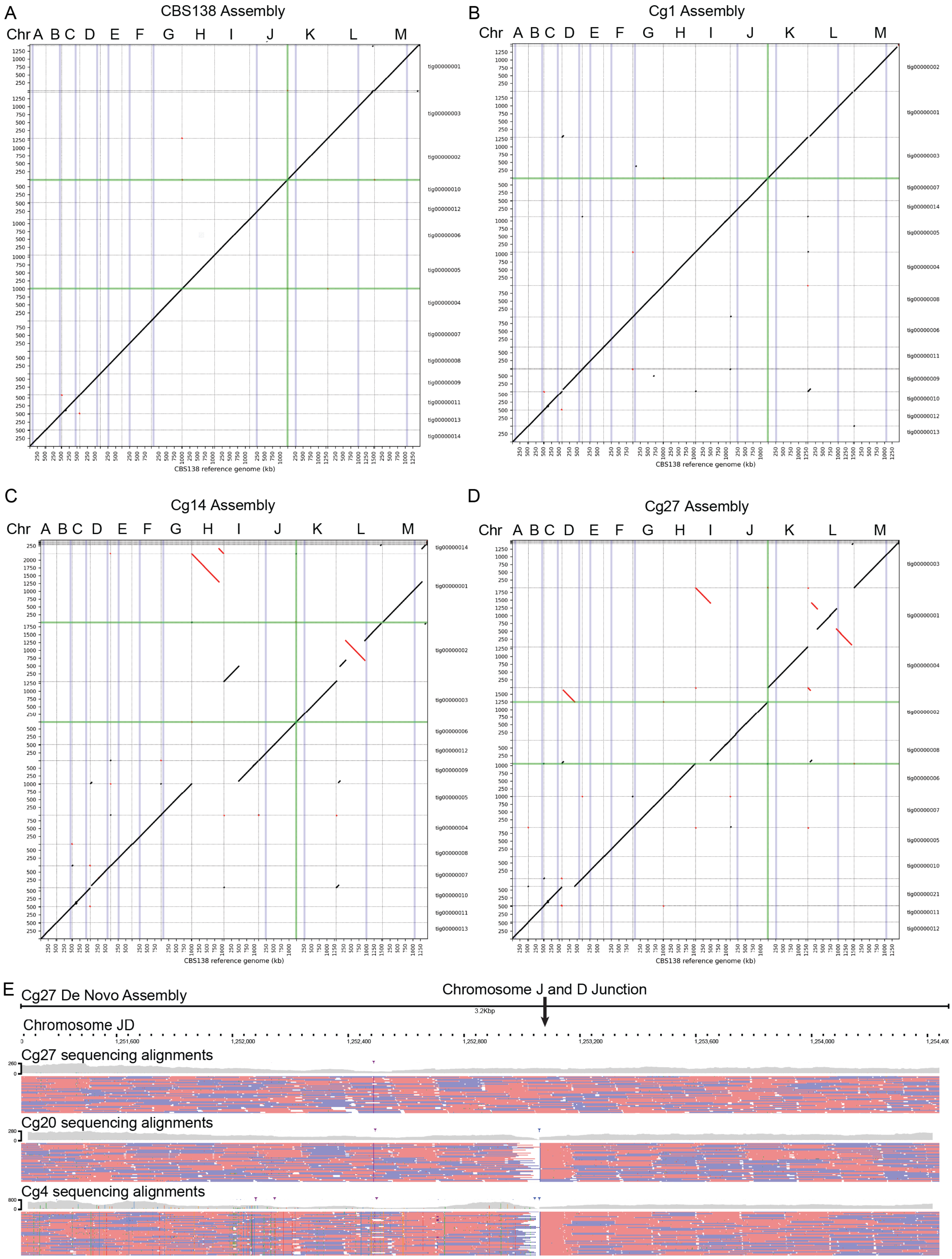
A
CBS138 Assembly
B
Cg1 Assembly
C
Cg14 Assembly
D
Cg27 Assembly
Chr A B C D E F G H I J K L M
CBS138 reference genome (kb)
E
Cg27 De Novo Assembly
Chromosome J and D Junction
3.2Kbp
Chromosome JD
1,251,600
1,252,000
1,252,400
1,252,800
1,253,200
1,253,600
1,254,000
Cg27 sequencing alignments
Cg20 sequencing alignments
Cg4 sequencing alignments

Fig. S5**.** ***C. glabrata* strain genomic structure.** (A-D) Dot plots comparing the genome structures of indicated strains to CBS138 reference genome. Black lines indicate alignment between contigs/chromosomes, red lines indicate inversions, and green cross sections indicate the location of *AWP11*. (E) Illumina sequencing alignments to Cg27 de novo genome assembly at the site of the JD chromosomal fusion. Red and blue bars represent individual reads and gray histogram represents the number of reads at each point.

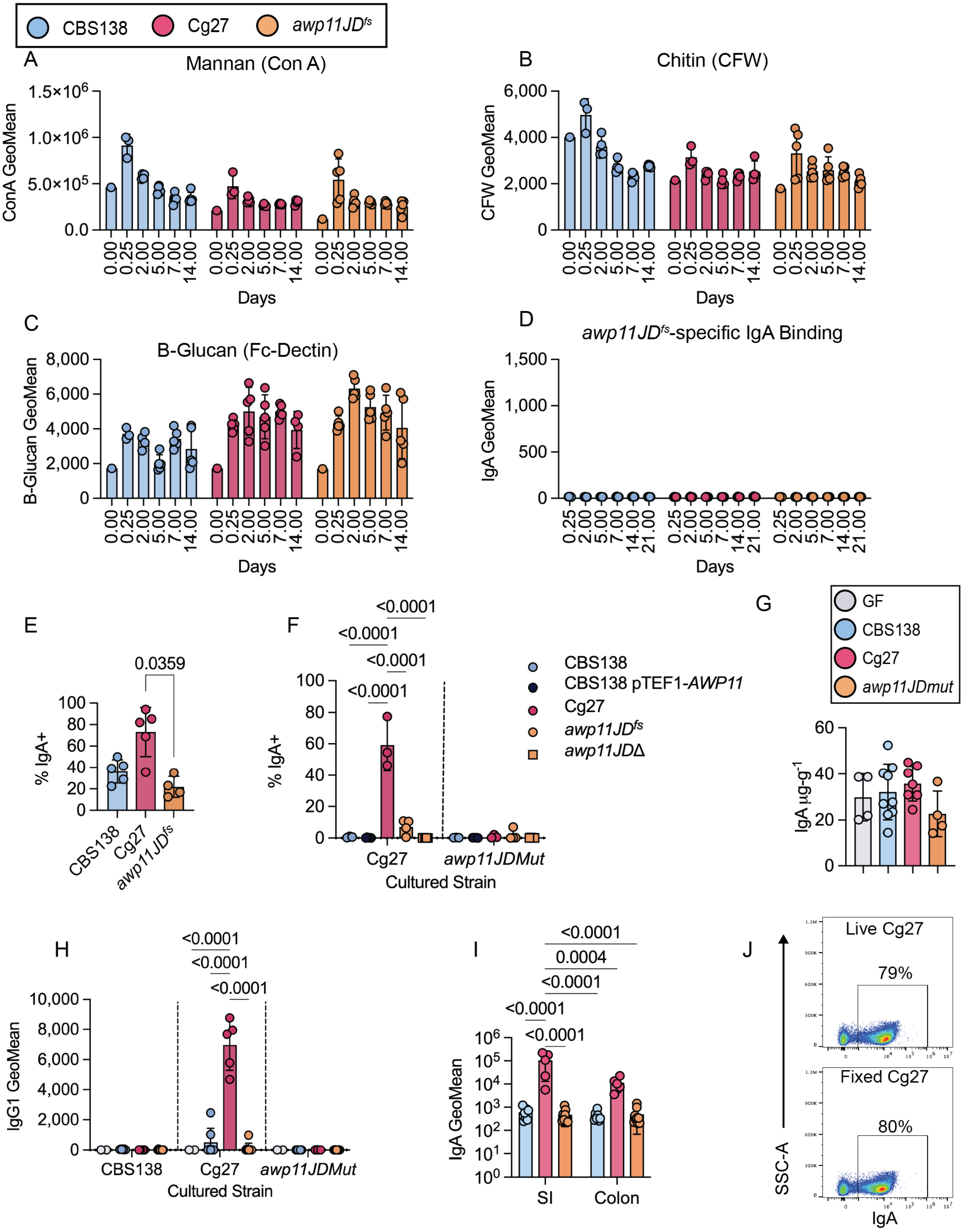

CBS138
Cg27
awp11JDfs
A
Mannan (Con A)
ConA GeoMean
Days
B
Chitin (CFW)
CFW GeoMean
C
B-Glucan (Fc-Dectin)
B-Glucan GeoMean
D
awp11JDfs-specific IgA Binding
IgA GeoMean
E
0.0359
% IgA+
F
<0.0001
CBS138 pTEF1-AWP11
awp11JDΔ
Cultured Strain
awp11JDMut
G
GF
awp11JDmut
IgA µg-g-1
H
IgG1 GeoMean
I
0.0004
SI
Colon
J
Live Cg27
79%
Fixed Cg27
80%
SSC-A
IgA

Fig. S6. **Awp11 is necessary for *C. glabrata*-specific IgA in vivo.** Time course of in vivo *C. glabrata* (A) mannan, (B) chitin, and (C) β-glucan cell wall components in stool measured by flow cytometry. Day 0 represents YPD grown *C. glabrata* cultures. (D) IgA geomean of cultured *awp11JDmut* incubated with paired stool wash from antibiotic treated mice. (E) Frequency of IgA binding in SI contents in vivo at day 21 from antibiotic treated mice. (F) IgA frequency of day 21 cleared intestinal content wash incubated with indicated cultured strains from antibiotic treated mice. n=3-5 mice per group from one independent experiment. (G) Total IgA in cleared intestinal wash (from SI) gnotobiotic mice. (H) IgG1 geomean from day 21 serum from incubated with indicated cultured strains. (I) In vivo *C. glabrata* IgA binding geomean in day 21 SI and colon content from monocolonized gnotobiotic mice. (J) Representative flow cytometry plots of IgA binding to live or fixed Cg27 following incubation with Cg27 monocolonized GF SI contents. All data are mean +/- SD. Each data point represents an individual mouse. (E, G) Kruskal-Wallis with Dunn's Multiple Comparison test, (F,H,I) Two-way ANOVA.

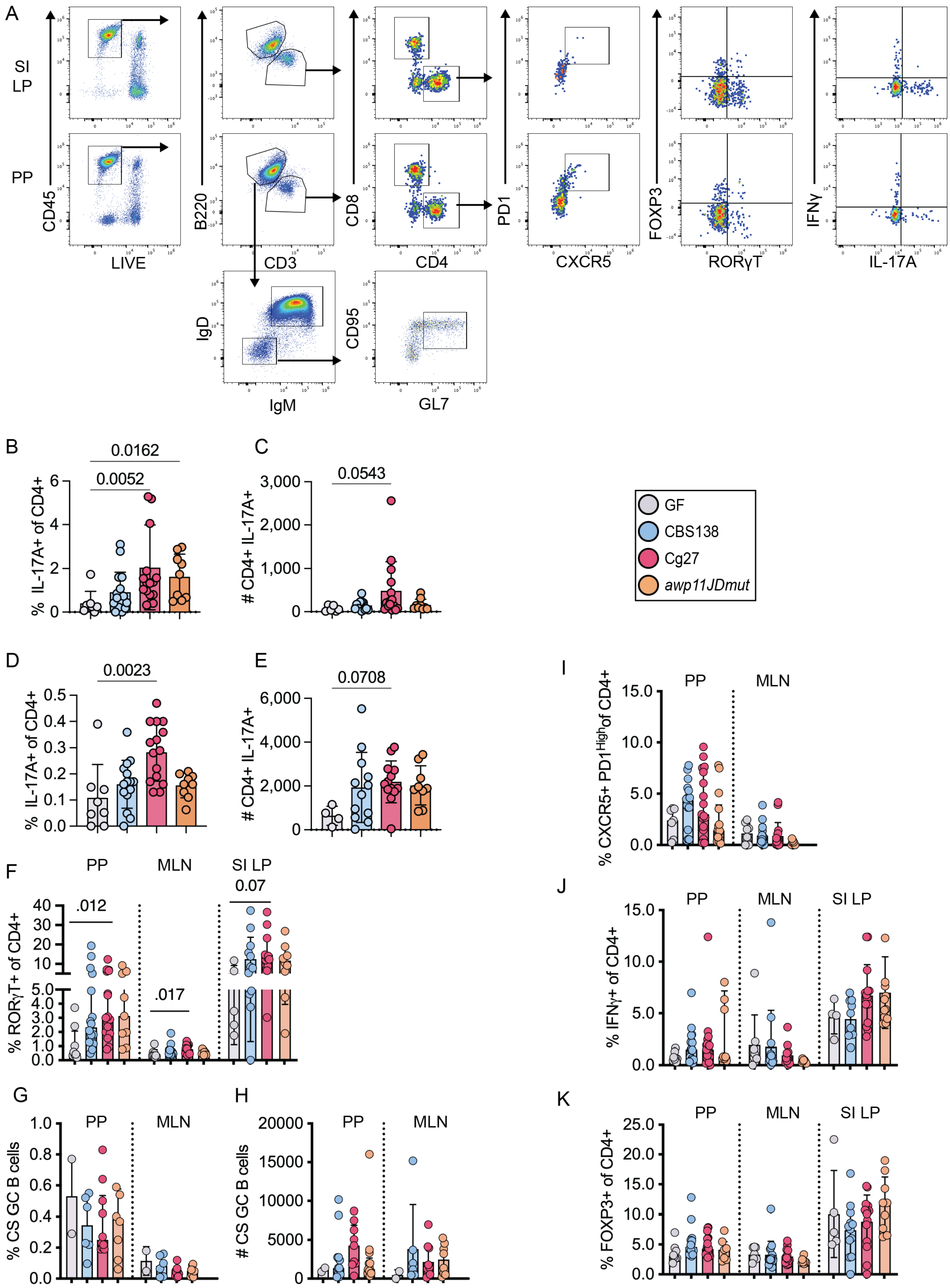
A
SI LP
PP
CD45
LIVE
B220
CD3
CD8
CD4
PD1
CXCR5
FOXP3
RORγT
IFNγ
IL-17A
IgD
IgM
CD95
GL7
B
% IL-17A+ of CD4+
0.0162
0.0052
C
# CD4+ IL-17A+
0.0543
GF
CBS138
Cg27
awp11JDmut
D
% IL-17A+ of CD4+
0.0023
E
# CD4+ IL-17A+
0.0708
I
% CXCR5+ PD1High of CD4+
PP
MLN
F
% RORγT+ of CD4+
PP
MLN
SI LP
.012
.017
0.07
J
% IFNγ+ of CD4+
PP
MLN
SI LP
G
% CS GC B cells
PP
MLN
H
### CS GC B cells
PP
MLN
K
% FOXP3+ of CD4+
PP
MLN
SI LP

Fig. S7. **Characterizing mucosal T cell responses in *C. glabrata* colonized gnotobiotic mice.** (A) Representative gating strategy from (Top) Peyer's patches and (Bottom) SI lamina propria of gnotobiotic mice colonized with *C. glabrata* clinical isolates. CD4+ T cells were delineated as Live CD45+ B220- CD3+ CD8a- cells and then gated on IL-17A+, IFNγ+, RORγT+ (Th17), FOXP3+ (Treg), or CXCR5+ PD1high (Tfh). Frequency (B,D) and total cell number (C,E) of CD4+ IL-17A+ cells in PP (B-C) and MLN (D-E). (F) Frequency of RORγT+ of CD4+ T cells in PP, MLN, and SI LP. Frequency (G) and total cell count (H) of GC B cells (Live CD45+ B220+ CD3- IgD- IgM- CD95+ GL7+) in PP and MLN. Frequency of (I) CXCR5+ PD1high (Tfh), (J) IFNγ+, and (K) FOXP3+ of CD4+ T cells in PP, MLN, and SI LP. n=4-13 mice per group, from 4 independent experiments. All data are mean +/- SD. Each data point represents an individual mouse. (B-K) Kruskal-Wallis with Dunn's Multiple Comparison test analyzed within tissues only.

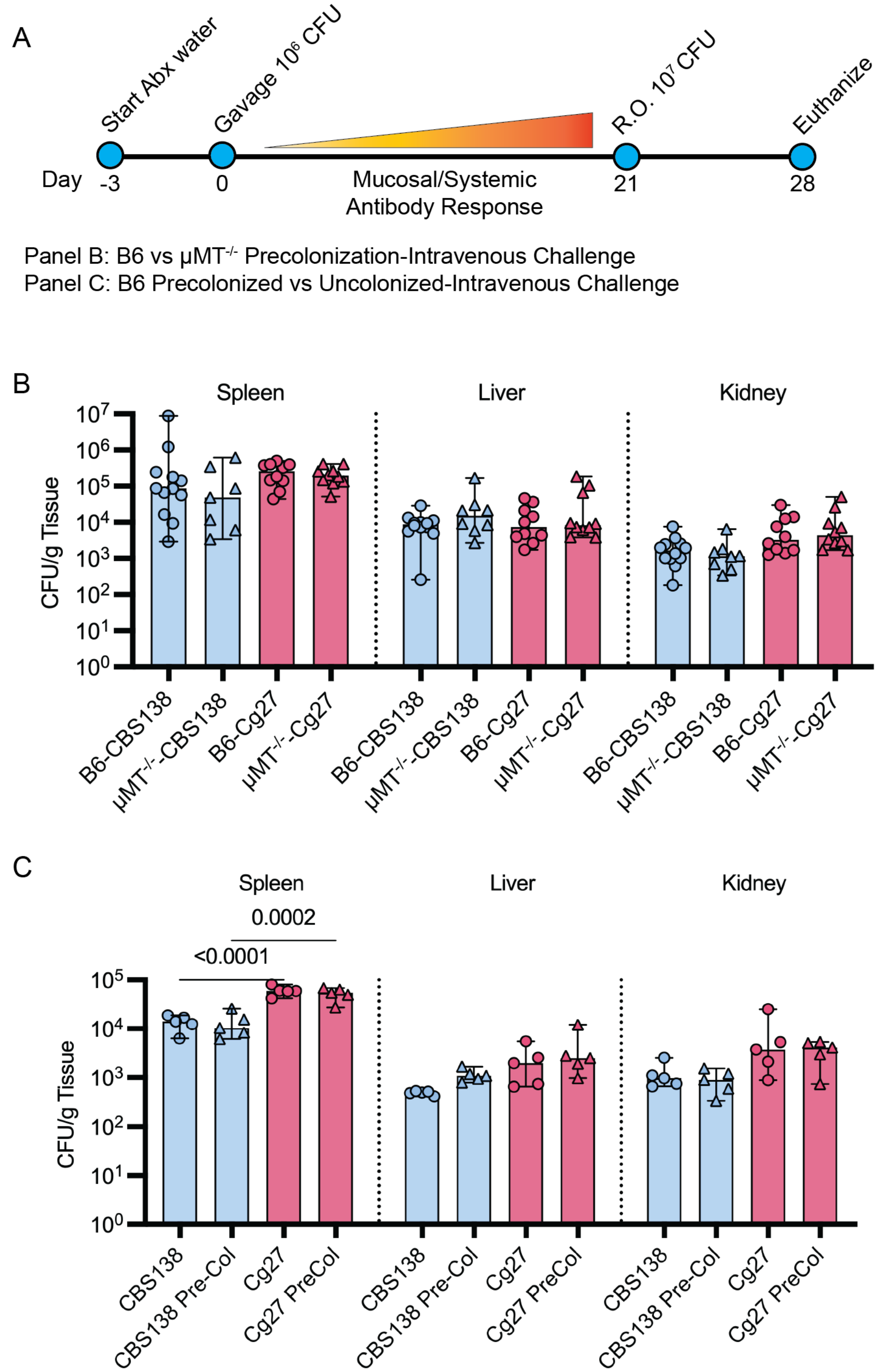


Fig. S8. **_C. glabrata_ Intestinal colonization-intravenous dissemination model.** (A) Timeline for *C. glabrata* intestinal colonization and dissemination model in B6 and μ$MT^{-/-}$ or B6 +/- intestinal colonization. (B-C) Fungal burden represented as CFU/g tissue in spleen, liver, and kidneys 7 days after intravenous infection with *C. glabrata* strains CBS138 or Cg27 (day 28). n=7-12 mice per group. All data are mean +/- SD. Each data point represents an individual mouse. Data analyzed by one-way ANOVA with Tukey's multiple comparison test within tissues only.

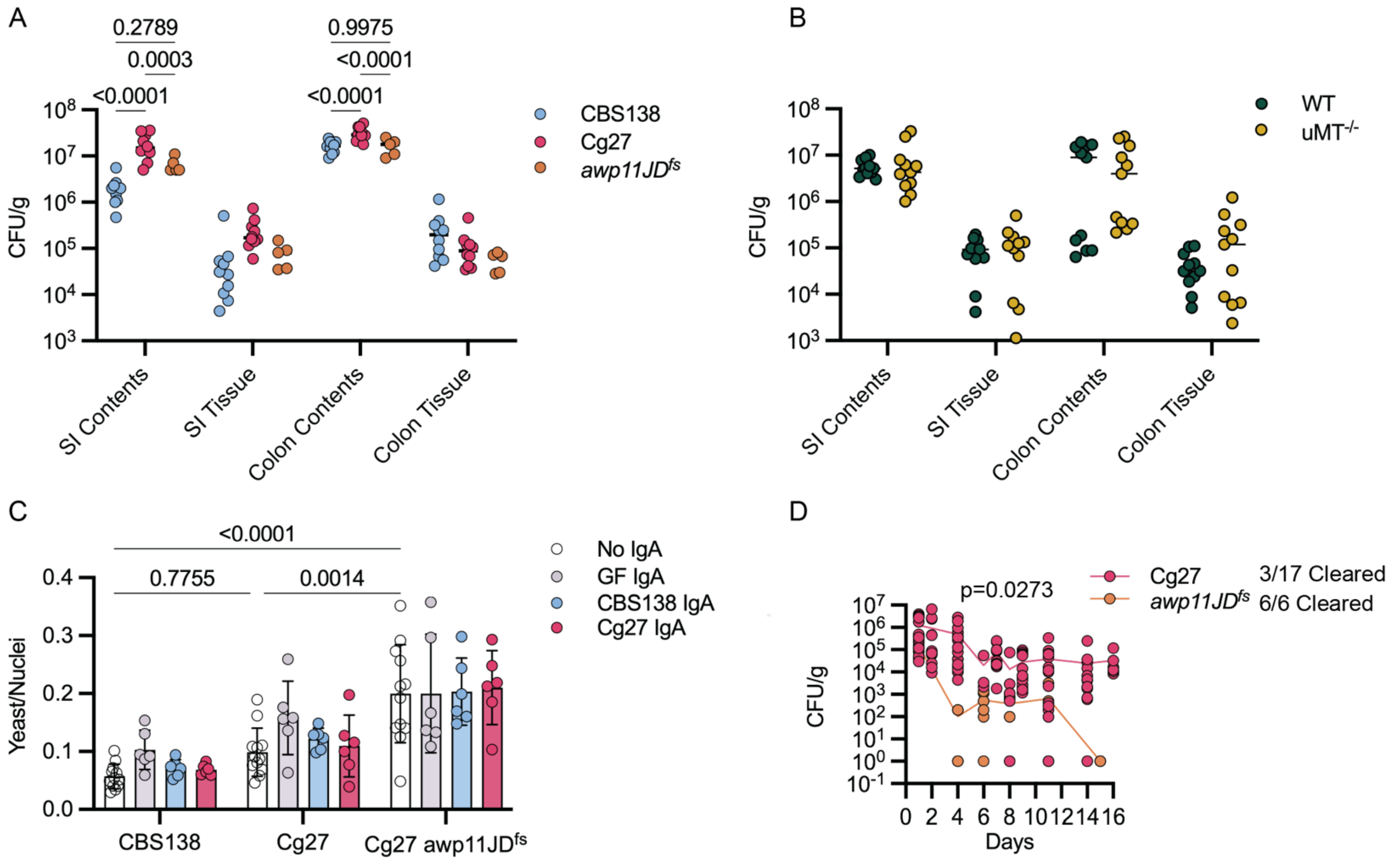


Fig. S9. **Awp11 does not promote adherence to colon epithelial cells, tissue association, nor short term colonization.** (A-B) Intestinal content and tissue fungal burden represented as CFU-gram$^{-1}$ contents/tissue. (C) *C. glabrata* cell count normalized to the colon epithelial (CACO-2) nuclei count per image. (D) Fecal fungal burden overtime represented as CFU/gram stool with ratio of mice who cleared *C. glabrata* from stool. (A-B) p values calculated by ordinary two-way ANOVA using Šídák's multiple comparisons test. n=11 mice per group across three independent experiments. (C) Two-way ANOVA with Tukey's multiple comparison test. n=6-12 replicates per group. Each dot represents the average of three images in one well across one experiment. (D) *Paired t-* test, n=6-17 mice per group across Data independent experiment. Each dot represents an individual mouse.

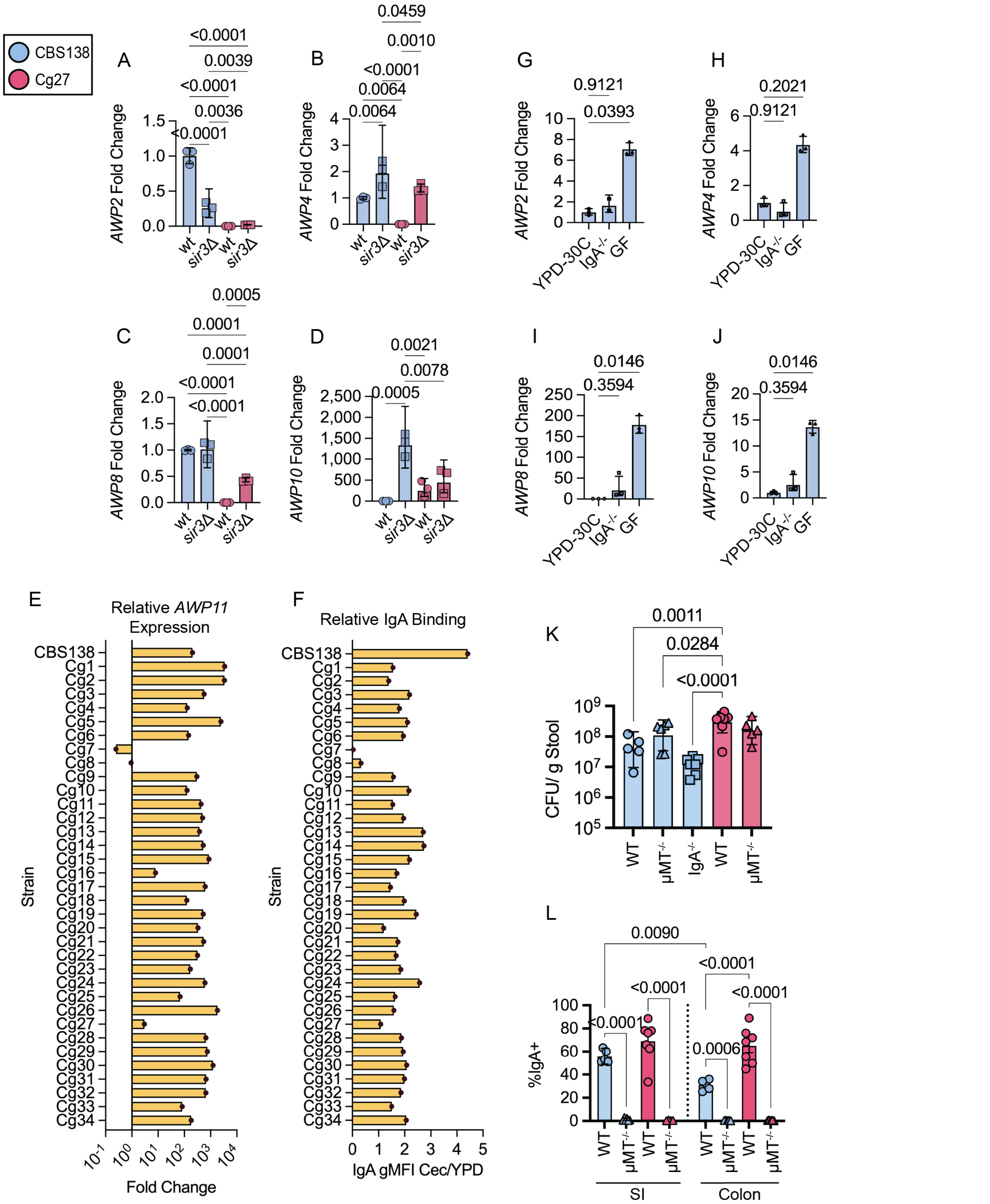


Fig. S10. **Intestinal environment promotes *AWP11* expression.** (A) *AWP2*, (B) *AWP4*, (C) *AWP8*, and (D) *AWP10* qRT-PCR analysis of WT and *sir3Δ* CBS138 and Cg27 grown in vitro at 30°C in YPD. (E-F) *C. glabrata* clinical isolates grown for 3 hours in YPD at 30°C or IgAKO mouse cecal contents at 37°C. n=3 from one independent experiment. (E) Cecal content *AWP11* expression measured by qRT-PCR relative to YPD expression in the same strain. Data represented as fold change normalized to an internal housekeeping gene, *ACT1*. (F) Relative IgA geomean of *C. glabrata* isolates following incubation with cecal wash supernatant from gnotobiotic mice colonized with Cg27 for 21 days. Data are represented as IgA geomean of cecal content sample divided by IgA geomean of YPD grown sample. CBS138 (G) *AWP2*, (H) *AWP4*, (I) *AWP8*, and (J) *AWP10* expression measured by qRT-PCR following incubation in YPD or IgAKO or GF cecal contents. Data represented as fold change normalized to an internal housekeeping gene, *ACT1* and YPD groups. n=3 from one independent experiment. (K) Fungal burden represented as CFU-$g^{-1}$ stool of CBS138 or Cg27 colonized WT, $\mu MT^{-/-}$, or IgAKO mice at day 21 post colonization. (L) Frequency of IgA binding in SI (left) or colon (right) contents in vivo at day 21 from antibiotic treated mice colonized with indicated strains. Data are mean +/- SD. Each data point represents an individual mouse. (A-B,G-L) One-way ANOVA with Tukey's multiple comparison test.
